# The activity-dependent transcription factor Npas4 regulates IQSEC3 expression in somatostatin interneurons to mediate anxiety-like behavior

**DOI:** 10.1101/659805

**Authors:** Seungjoon Kim, Dongseok Park, Jinhu Kim, Dongsoo Lee, Dongwook Kim, Hyeonho Kim, Sookyung Hong, Jongcheol Jeon, Jaehoon Kim, Eunji Cheong, Ji Won Um, Jaewon Ko

## Abstract

Organization of mammalian inhibitory synapses is thought to be crucial for normal brain functions, but the underlying molecular mechanisms have been still incompletely understood. IQSEC3 (IQ motif and Sec7 domain 3) is a guanine nucleotide exchange factor for ADP-ribosylation factor (ARF-GEF) that directly interacts with gephyrin. Here, we show that GABAergic synapse-specific transcription factor, Npas4 (neuronal PAS domain protein 4) directly binds to the promoter of *Iqsec3* and regulates its transcription. Strikingly, an enriched environment (EE) induced Npas4 upregulation and concurrently increased IQSEC3 protein levels specifically in mouse CA1 *stratum oriens* layer somatostatin (SST)-expressing GABAergic interneurons, which are compromised in *Npas4*-knockout (KO) mice. Moreover, expression of wild-type (WT) IQSEC3, but not a dominant-negative (DN) ARF-GEF–inactive mutant, rescued the decreased GABAergic synaptic transmission in Npas4-deficient SST interneurons. Concurrently, expression of IQSEC3 WT normalized the altered GABAergic synaptic transmission in dendrites, but not soma, of Npas4-deficient CA1 pyramidal neurons. Furthermore, expression of IQSEC3 WT, but not IQSEC3 DN, in SST-expressing interneurons in CA1 SST *Npas4*-KO mice rescued the altered anxiety-like behavior. Collectively, our results suggest that IQSEC3 is a key GABAergic synapse component that is directed by Npas4 activity- and ARF activity-dependent gene programs in SST-expressing interneurons to orchestrate the functional excitation-to-inhibition balance.

## Introduction

An essential function of the nervous system is to generate optimal behavioral responses to diverse environmental information; moreover, combinatorial interactions between genetic components and sensory cues are essential for driving discrete steps in brain development^1,2,3,4^. Neuronal activity and subsequent calcium influx activate specific signaling cascades that cause nuclear translocation of various transcription factors, resulting in the rapid induction of an early-response gene-expression program that, in turn, induces transcription of late-response genes that locally regulate synaptic development and plasticity. This paradigm has been well established at both glutamatergic-producing and γ-aminobutyric acid-producing (GABAergic) synapses^3, 5^. Transcription factors that are commonly induced by synaptic activity regulate distinct sets of target genes to control the strength, location, and number of both glutamatergic and GABAergic inputs^6^. However, which specific transcription factors couple with key synapse-organizing proteins to orchestrate the regulation of synapse development at both transcriptional and post-transcriptional levels remains incompletely understood^3^.

One prominent transcription factor that specifically controls GABAergic synapse development is Npas4 (neuronal PAS domain protein 4)^7,8,9,10,11, 12^, which is required for neuronal excitability in the adult dentate gyrus (DG) and for new and reactivated fear memories^13, 14^. Npas4 recruits RNA polymerase II to promoters and enhancers of target genes that are regulated by neuronal activity in CA3 hippocampal subfields, thereby contributing to short- and long-term contextual memory^15^. Deletion of *Npas4* abolishes depolarization-induced mRNA expression of immediate-early genes, such as Arc/Arg3.1, c-Fos, Zif268/Egr1, and brain-derived neurotrophic factor (BDNF)^15^. Importantly, Npas4 expression levels dictate the degree of inhibition in specific neuronal compartments by organizing distinct neuron-type– specific genetic programs^8, 9^. Notably, BDNF regulates somatic, but not dendritic, inhibition in neural circuits of CA1 pyramidal neurons, and is responsive only to increased inhibitory inputs in glutamatergic neurons^8, 9^. In addition, Npas4 mediates experience-dependent spine development in olfactory bulb interneurons (INs) through control of Mdm2 expression, confers neuroprotection against kainic acid-induced excitotoxicity in hippocampal neurons through control of synaptotagmin-10 (Sy10) expression, and organizes the structure and strength of hippocampal mossy fiber-CA3 synapses during contextual memory formation through control of polo-like kinase 2 (Plk2) expression^16,17,18^. Moreover, in response to heightened neuronal activity, Npas4 is recruited to activity-dependent regulatory elements by ARNT2, which is normally complexed with NCoR2, a co-repressor of neuronal activity-regulated gene expression under basal states^19^. These prior studies suggest that synaptic roles of Npas4 vary in brain region-, synaptic-, and cellular context-dependent manner. Despite these insights, the sheer number of Npas4 target genes has hampered efforts to fully understand the mechanisms underlying Npas4-mediated GABAergic synapse development. IQSEC3 (IQ motif and Sec7 domain 3; also known as BRAG3 or SynArfGEF) was previously isolated as a putative Npas4 target^8^. However, direct functional coupling of Npas4 with IQSEC3 in the context of GABAergic synapse development *in vivo* has not been investigated.

Postsynaptic scaffolding proteins organize functional synapses, provide platforms for postsynaptic receptors, and regulate downstream signaling cascades^20, 21^. Arguably the most extensively studied scaffolding protein at GABAergic synapses is gephyrin^22, 23^. We recently reported that gephyrin directly interacts with IQSEC3 to promote GABAergic synapse formation^24, 25^. IQSEC3, a member of the brefeldin A-resistant ADP ribosylation factor guanine exchange factor (ARF-GEF) family, is exclusively localized at GABAergic synapses^24,25,26,27^. IQSEC3 overexpression increases the number of postsynaptic gephyrin and presynaptic GAD67 puncta, whereas IQSEC3 knockdown (KD) specifically decreases gephyrin cluster size in cultured hippocampal neurons, without altering gephyrin puncta density^24^. In addition, IQSEC3 KD in the dentate gyrus induced severe seizure activity, an effect that was reversed by restricted expression of somatostatin (SST) peptides in SST^+^ neurons^28^, suggesting that IQSEC3 may be critical for maintenance of network activities *in vivo*. However, the brain-region specific, neuronal activity-dependent role of IQSEC3 *in vivo* has yet to be identified.

Here, we provide the first evidence demonstrating that IQSEC3 mediates activity-dependent GABAergic synapse development, coupled with Npas4-dependent transcriptional programs, specifically in SST-expressing interneurons. Npas4 binds to the promoter of *Iqsec3* to regulate *Iqsec3* expression levels in a manner that depends on synaptic activity, which concurrently increase Npas4 levels. Remarkably, EE-induced upregulation of IQSEC3 proteins is prominent in SST-expressing interneurons in the hippocampal CA1 stratum oriens layer, suggesting a unique role for IQSEC3 in shaping hippocampal circuitries. In keeping with this observation, IQSEC3 regulates Npas4-dependent spontaneous GABAergic synaptic transmission in SST-expressing interneurons, and further modulates the mode of Npas4-mediated GABAergic synaptic transmission in apical dendrites (but not soma) of CA1 pyramidal neurons, a function that also requires its ARF-GEF activity. Furthermore, expression of IQSEC3 WT, but not an ARF-GEF activity-deficient IQSEC3 mutant, in SST-expressing interneurons restored the altered anxiety-like behaviors observed in conditional knockout (KO) mice lacking *Npas4* in SST interneurons. We further suggest that IQSEC3 is a downstream factor for Npas4 in sculpting specific neuronal inhibition in distal dendritic domains by providing inhibitory inputs onto excitatory neurons, hinting at its crucial roles in shaping activity-dependent specific inhibitory neural circuits and controlling a selective set of mouse behaviors.

## Results

### Npas4 binds to the *IQSEC3* promoter

The structural and functional plasticity of GABAergic synapses are dynamically modulated in a neuronal activity-dependent manner^29, 30^. We thus hypothesized that IQSEC3 might be involved in activity-dependent GABAergic synapse development. Among GABAergic synapse-specific transcription factors, Npas4 appears to be a prominent candidate factor with the potential to play this role^7, 31^. Npas4 orchestrates the expression of a wide variety of target genes, notably including that encoding BDNF, which regulates GABAergic synapse development to mediate domain-specific inhibition in particular cell-types^7,8, 9^. Intriguingly, deep sequencing of Npas4-bound DNA revealed IQSEC3 as a putative Npas4 target^8^, and a recent genome-wide ChIP-seq analysis showed strong enrichment of Npas4 at the upstream region near the transcription start site of the *IQSEC3* gene^32^ (**Figure 1a**). Intriguingly, we noted that this Npas4 peak contains a single Npas4 consensus CACG motif (**Figure 1a**). Thus, we tested whether IQSEC3, like BDNF, could also be an Npas4 target that acts to promote GABAergic synapse development. To this end, we cotransfected HEK293T cells with a *Iqsec3* promoter-luciferase reporter construct and a vector expressing Npas4. Ectopic expression of Npas4 increased luciferase expression ∼3-fold; notably, this increase was abolished by mutation of the Npas4 consensus site in the *Iqsec3* promoter region (**Figure 1b**). We further performed luciferase reporter assays in cultured cortical neurons transfected with Npas4 wild-type (WT) or control, and treated with 55 mM KCl to trigger membrane depolarization (**Figure 1c**). We found that Npas4 WT similarly increased luciferase expression of IQSEC3 by ∼4-fold (**Figure 1c**). Electrophoretic mobility shift assays (EMSAs) confirmed these results, showing that a radiolabeled *Iqsec3* WT oligoduplex probe containing purified Npas4 protein yielded a single bound complex; consistent with promoter-reporter assays, this DNA-protein interaction was abolished by mutation of the Npas4 consensus site (**Figure 1d**). Chromatin immunoprecipitation (ChIP) assays also demonstrated that Npas4 binds to the *Iqsec3* promoter in primary hippocampal cultured neurons treated with 55 mM KCl to trigger membrane depolarization (**Figure 1e**). No Npas4-binding was detected in untreated hippocampal cultured neurons (**Figure 1e**). These data suggest that Npas4 can function as a DNA-binding transcriptional activator of *IQSEC3*.

**Fig. 1.**
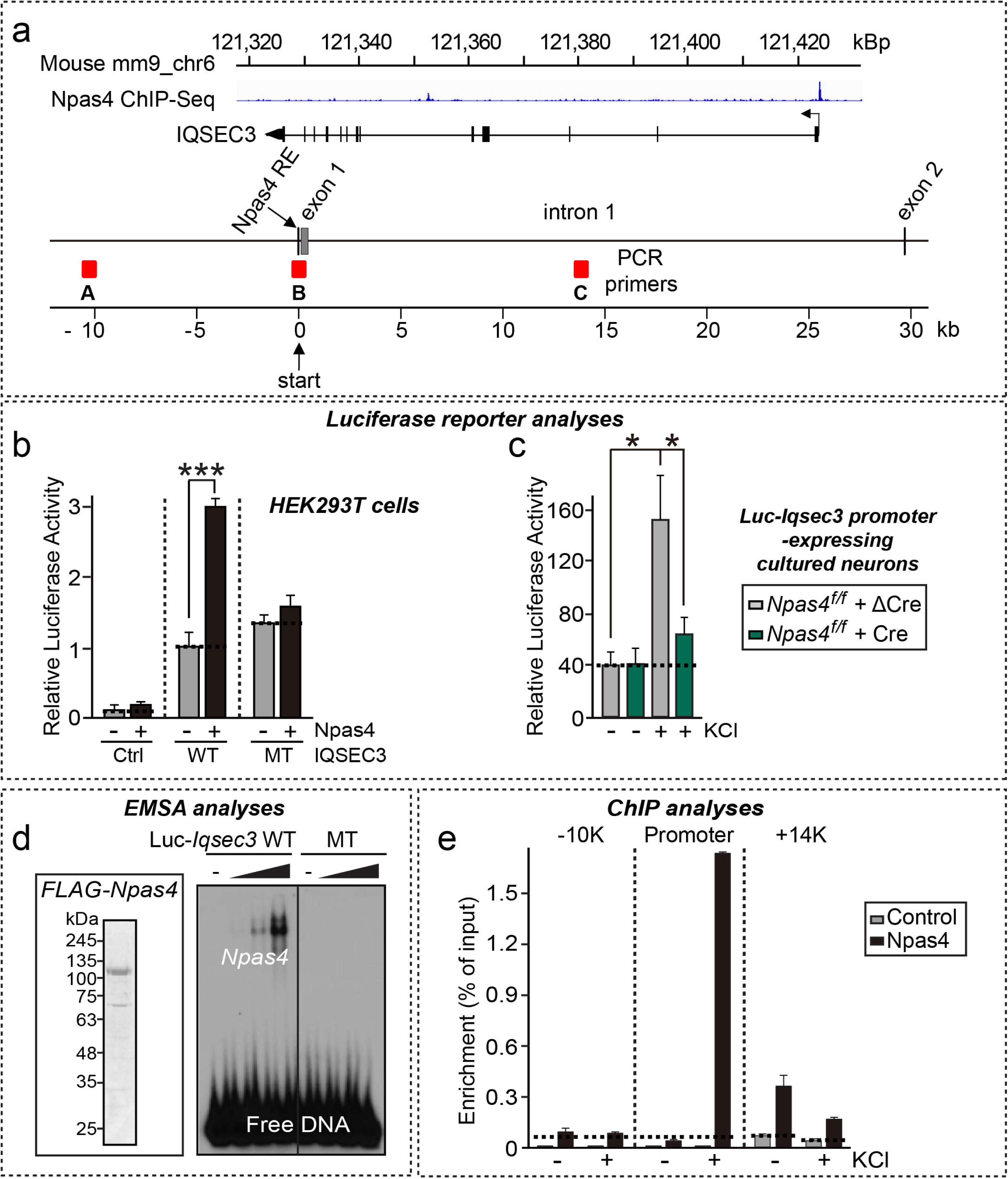
Npas4 binds to the IQSEC3 promoter in an activity-dependent manner. **a** Schematic representation of the Iqsec3 locus indicating the three amplicons used for RT-qPCR. An Npas4-binding site identified by ChIP-seq analysis^32^ is indicated as an Npas4 responsive element (Npas4 RE). **b** HEK293T cells were cotransfected with a luciferase reporter construct (100 ng) containing a *Iqsec3* proximal promoter region (–280 to +36) together with a vector expressing Npas4 (40 ng), as indicated. MT indicates a promoter-reporter construct harboring a mutation in the consensus Npas4-binding sequence. Luciferase activity in cell extracts was assayed and expressed as relative to luciferase activity in cells transfected with a luciferase-reporter construct containing the WT *Iqsec3* promoter region (defined as 1). Values are expressed as means ± SEM from three independent experiments (\*\*\**p* < 0.001 vs. Mock-transfected controls; Mann-Whitney U Test). **c** Cortical cultured neurons derived from Npas4*^f/f^* mice were infected with lentiviruses expressing ΔCre or Cre at DIV3, and transfected with a luciferase reporter construct containing an *Iqsec3* proximal promoter region at DIV10. Luciferase activity in extracts of neurons left untreated or treated with 55 mM KCl for 2 hours was assayed at DIV14. Values are expressed as means ± SEM from three independent experiments (\**p* < 0.05; non-parametric ANOVA with Kruskal-Wallis test, followed by *post hoc* Dunn’s multiple comparison test.). **d** EMSAs employing purified FLAG-tagged Npas4 protein (10–90 ng) and a probe corresponding to nucleotide −110 to −69 within the *Iqsec3* proximal promoter are shown. An inset shows an image of a Coomassie Blue-stained SDS-PAGE gel of the purified FLAG-Npas4 protein used in experiments. **e** ChIP-seq binding profiles for Npas4 at the *Iqsec3* locus in KCl-treated mouse cortical neurons. Data were downloaded from the GEO website (http://www.ncbi.nlm.nih.gov/geo); GEO accession number, GSE21161^31^. The total number of tags was normalized to the corresponding input.

### IQSEC3 functions as an Npas4 target to promote GABAergic synapse development

To further determine whether IQSEC3, analogous to BDNF^7^, is involved in Npas4 actions on GABAergic synapse development, we infected cultured neurons with lentiviruses expressing shRNAs against *Iqsec3*, *Bdnf*, or control shRNA at 3 days *in vitro* (DIV3); transfected them at DIV10 with EGFP alone, or EGFP and myc-Npas4; and then immunostained them at DIV14 for gephyrin. Overexpression of Npas4 significantly increased gephyrin puncta density in both dendrites and soma, an effect that was completely abolished by IQSEC3 KD (**Supplementary Figure 1**). These effects were distinct from the previously reported BDNF-KD effects on an Npas4 minigene, which led to altered GABAergic synaptic transmission specifically in the soma, but not in the dendritic compartment^8^ (**Supplementary Figure 1**).

These data reinforce the conclusion that IQSEC3 acts downstream of Npas4 to regulate activity-dependent GABAergic synapse development in cultured hippocampal neurons.

### IQSEC3 expression is specifically upregulated in somatostatin-positive interneurons of hippocampal CA1 region

It was previously reported that expression of *Npas4* and *Bdnf* mRNAs changes in response to neuronal depolarization^7, 8^. Thus, we asked whether *Iqsec3* mRNA expression is also responsive to changes in neuronal activity (**Figure 2** and **Supplementary Figure 2**). To test this, we manipulated the sensory experiences of juvenile wild-type (WT) mice by housing littermates in an enriched environment (EE), consisting of a running wheel and several novel objects that were regularly refreshed (**Figure 2a**, see ‘**Methods**’ for details). After 5 days in an EE setting, hippocampi were removed, sectioned, and immunostained for c-Fos as an indirect marker of neuronal activity. As expected, EE exposure drove robust c-Fos expression in the hippocampal CA1 region (**Figure 2b**). We then performed immunohistochemical analyses on sections from age-matched mice housed in a standard environment (SE) as a control using previously validated antibodies recognizing Npas4 and IQSEC3^7, 27^. Basal levels of IQSEC3 in the hippocampal CA1 region from mice housed under SE conditions were low (**Figure 2c, 2d**).

**Fig. 2.**
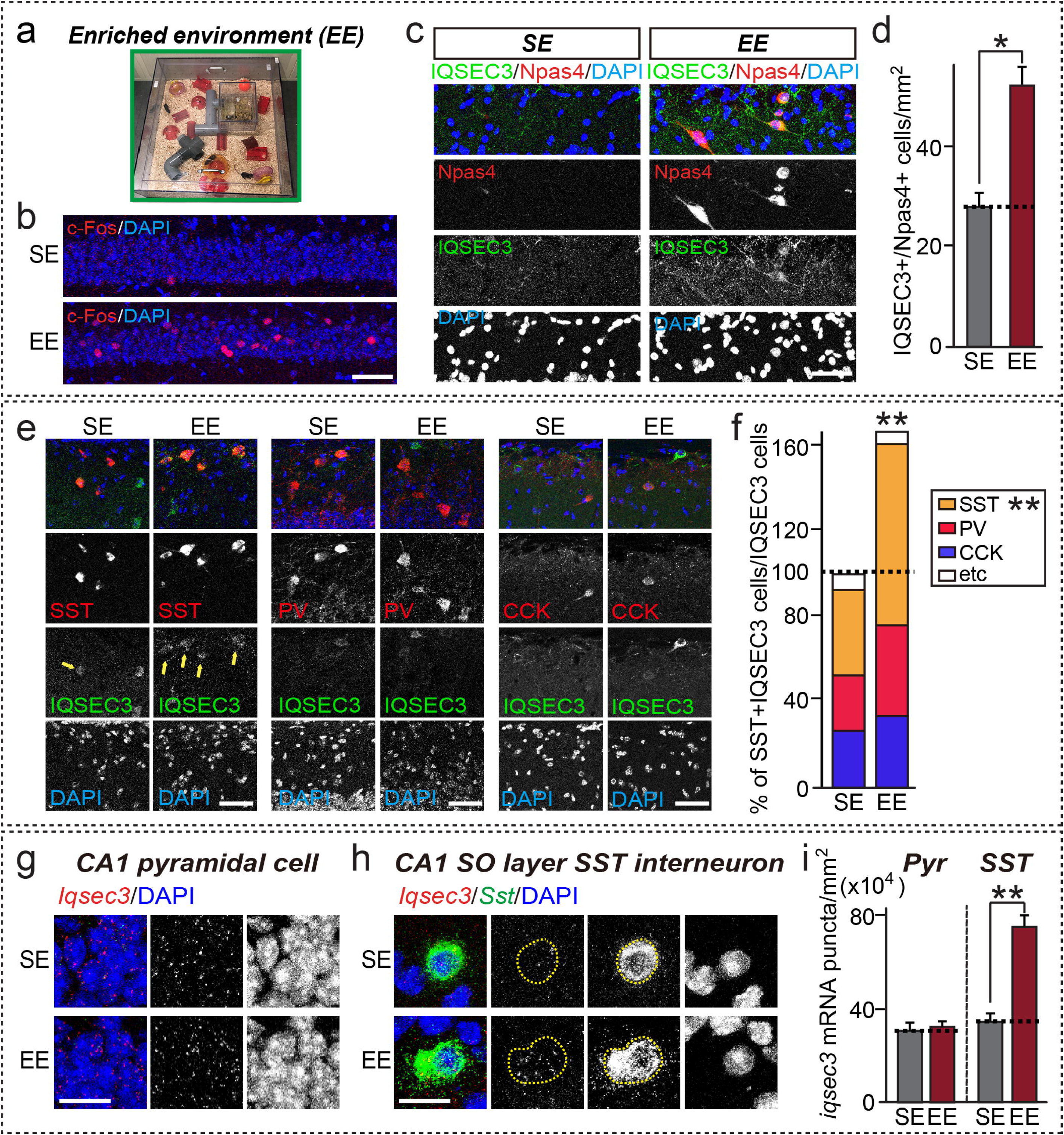
IQSEC3 protein level is regulated by neuronal activity *in vivo*. **a** The enriched environment (EE) setting, including tunnels, shelters, toys and running wheels for voluntary exercise, used in the current study. **b** Representative immunofluorescence images of hippocampal CA1 regions from adult (11-week-old) WT mice in a standard environment (SE) or EE setting were immunostained for c-Fos (red) and counterstained with DAPI (blue). **c** Representative immunofluorescence images of the hippocampal CA1 SO layer from WT mice in an SE or EE setting were immunostained for Npas4 (red) and IQSEC3 (green), and counterstained with DAPI (blue). **d** Bar graphs summarizing the density of IQSEC3/Npas4 double-positive neurons shown in (**c**). Data are presented as means ± SEMs (\**p* < 0.05; Mann-Whitney *U* test; n = 24 sections/4 mice for all conditions). **e** Representative immunofluorescence images of the hippocampal CA1 SO layer from WT mice in an SE or EE setting, immunostained for IQSEC3 (green), DAPI (blue), and SST (red; **left**), PV (red; **middle**), or CCK (red; **right**). Yellow arrows indicate SST-positive interneurons with upregulated IQSEC3 levels. Scale bar, 50 μm (applies to all images). **f** Bar graphs summarizing quantitative results shown in (**e**). Data are presented as means ± SEMs (\*\**p* < 0.01; Mann-Whitney *U* test; n = 18 − 24 sections/3 − mice). **g, h** Expression of *Iqsec3* (red) and *Sst* (green) mRNAs in adult WT mice in an SE or EE setting was visualized by *in situ* hybridization using RNAscope technology. Nuclei were stained with DAPI (blue); representative images of hippocampal CA1 neurons from the pyramidal (**g**) and SO layer (**h**) from mice reared under the indicated conditions are shown. Scale bar, 20 μ (applies to all images). **i** Quantitative analyses of RNAscope data measuring *Iqsec3* mRNA expression in pyramidal neurons or SST^+^ interneurons. Data are presented as means ± SEMs (\*\**p* < 0.01, nonparametric one-way ANOVA with Kruskal-Wallis test followed by *post hoc* Dunn’s multiple comparison test; n = 3-5 mice).

There was a significant increase in the number of Npas4^+^ neurons in CA1 *stratum oriens* (SO) layers from mice allowed to explore an EE relative to those maintained in an SE (**Figure 2c**). Notably, IQSEC3 expression was also concomitantly increased in Npas4-positive neurons, an effect that was prominent in SO layers of the CA1 hippocampus (**Figure 2c, 2d**). EE-induced upregulation of IQSEC3 in the SO layer of the CA1 hippocampus was prominently observed in SST, but not parvalbumin (PV) or cholecystokinin (CCK), GABAergic interneurons (**Figure 2e, 2f**). RNAscope-based *in situ* hybridization analyses also showed that EE induced marked increases in the number of SST^+^ neurons in the SO layer, accompanied by significant increases in IQSEC3 mRNA expression, without affecting the number of pyramidal neurons in the hippocampal CA1 region (**Figure 2e–g**), suggesting the operation of a specific transcriptional mechanism for IQSEC3 mRNAs selectively in SST^+^ neurons.

To determine whether an activity-dependent increase in Npas4 level is responsible for selective upregulation of IQSEC3 in the SO layer of the hippocampal CA1 region, we expressed either active Cre-combinase fused to EGFP (Cre) or inactive Cre-recombinase fused to EGFP (ΔCre, as a control) in floxed Npas4 mice (Npas4*^f/f^)* via adeno-associated virus (AAV)-mediated stereotactic injection, and housed the resulting mice in either SE or EE conditions (**Figure 3a**). Quantitative immunohistochemical analyses showed that EE induced a significant increase in the number of Npas4^+^ or IQSEC3^+^ neurons and Npas4 deletion blunted the this effect, reducing the number of IQSEC3^+^ neurons to a level comparable to that observed in SE-housed *Npas4*-cKO mice (**Figure 3b–e**). This blunting of the number of IQSEC3^+^ neurons was also recapitulated by specific deletion of Npas4 in SST^+^ neurons (**Figure 3f, 3g**; see **Supplementary Figure 2** for validation of the EE-induced upregulation of Npas4 level in Npas4-deficient SST^+^ brain sections), further supporting the idea that Npas4 induces upregulation of IQSEC3 specifically in SST^+^ neurons.

**Fig. 3.**
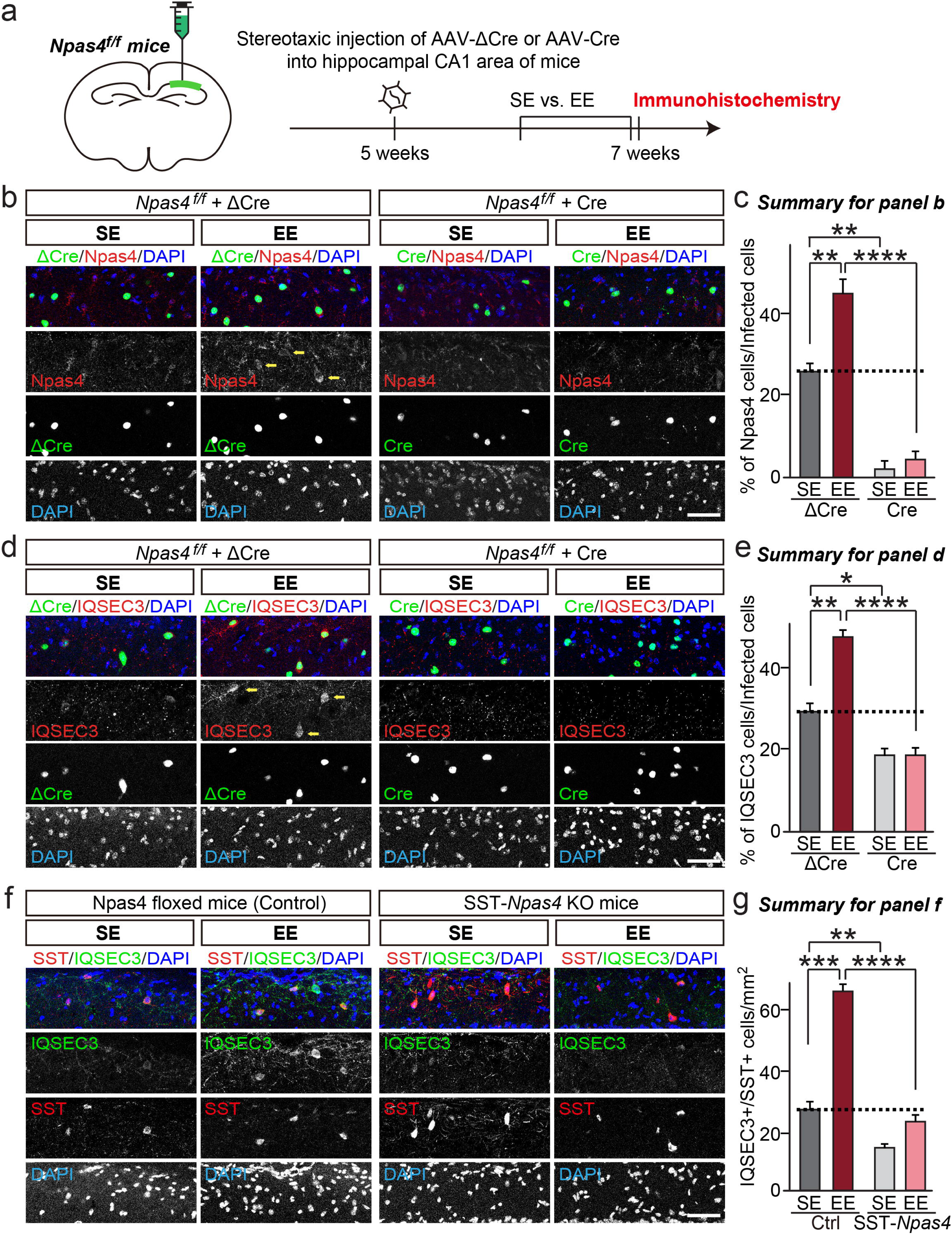
Upregulation of IQSEC3 protein level *in vivo* depends on the presence of Npas4. **a** Experimental scheme for immunohistochemistry. The CA1 region of the hippocampus of ∼9-week-old *Npas4*^f/f^ mice was bilaterally injected with AAVs expressing ΔCre or Cre. Mice were housed for 5 days in an SE or EE setting 9 days after AAV injections, and then analyzed by immunohistochemistry. **b, d** Representative immunofluorescence images of the hippocampal CA1 SO layer from *Npas4*^f/f^ mice was bilaterally injected with AAVs expressing ΔCre or Cre in an SE or EE setting, immunostained for Npas4 (red) (**b**) or IQSEC3 (red) (**d**) and DAPI (blue). Green fluorescence indicates neurons infected with AAV-ΔCre or AAV-ΔCre. Scale bar, 50 μ (applies to all images). **c, e** Bar graphs summarizing the percentage of Npas4^+^ (**c**) or IQSEC3^+^ (**e**) cells shown in (**b** and **d**). Data are presented as means ± SEMs (\**p* < 0.05; \*\**p* < 0.01, \*\*\*\**p* < 0.0001; non-parametric ANOVA with Kruskal-Wallis test, followed by *post hoc* Dunn’s multiple comparison test). **f,** Representative immunofluorescence images of the hippocampal CA1 SO layer from Control or SST-*Npas4* mice under SE or EE conditions, immunostained for SST (red), IQSEC3 (green), and DAPI (blue). Scale bar, 50 μm (applies to all images). **g,** Bar graphs summarizing the number of IQSEC3^+^ cells expressed in SST^+^ interneurons shown in (**f**). Data are presented as means ± SEMs (\*\**p* < 0.01, \*\*\**p* < 0.001, \*\*\*\**p* < 0.0001; non-parametric ANOVA with Kruskal-Wallis test, followed by *post hoc* Dunn’s multiple comparison test).

To further investigate the expression profile of IQSEC3 under pathological conditions in which network activity is severely heightened, we performed immunohistochemistry using ∼11 week-old adult mice that were either injected with saline (SA) or 15 mg/kg of KA (**Supplementary Figure 3a**). KA injections significantly increased Npas4^+^ cell numbers in both SP and SO layers of the hippocampal CA1 region, GCL and hilus of the DG (**Supplementary Figure 3b**). Strikingly, KA injections resulted in an increase in the number of IQSEC3^+^ cells, specifically in the SO layer, but not in the SP layer, molecular layer, or GCL of the DG (**Supplementary Figure 3b**). In addition, quantitative analyses using KA-injected Npas4 cKO mice showed that Npas4 deletions decreased the number of IQSEC3^+^ cells to a level comparable to that observed in SA-injected WT mice (**Supplementary Figure 3c, 3d**). Furthermore, the KA injection-dependent upregulation of IQSEC3 in the SO layer of the CA1 hippocampus was prominently observed in SST, but not PV or CCK, GABAergic interneurons (**Supplementary Figure 3e, 3f**). Collectively, these data show that IQSEC3 levels are preferentially altered specifically in SST interneurons following induction of neuronal activity *in vivo*.

### IQSEC3 is required for postsynaptic GABAergic synaptic transmission in Npas4-deficient SST-expressing interneurons

To determine whether Npas4-induced increases in IQSEC3 levels in SST-expressing interneurons has an impact on postsynaptic GABAergic transmission, we crossed mice carrying the Cre-driver SST-Cre^33^ and the Ai9 reporter mouse line^34^ with Npas4 floxed mice (Npas4*^f/f^* mice), and exposed the resulting mice to EE conditions to induce Npas4 protein expression, as previously described^8, 35^ (**Figure 4a**).

**Fig. 4.**
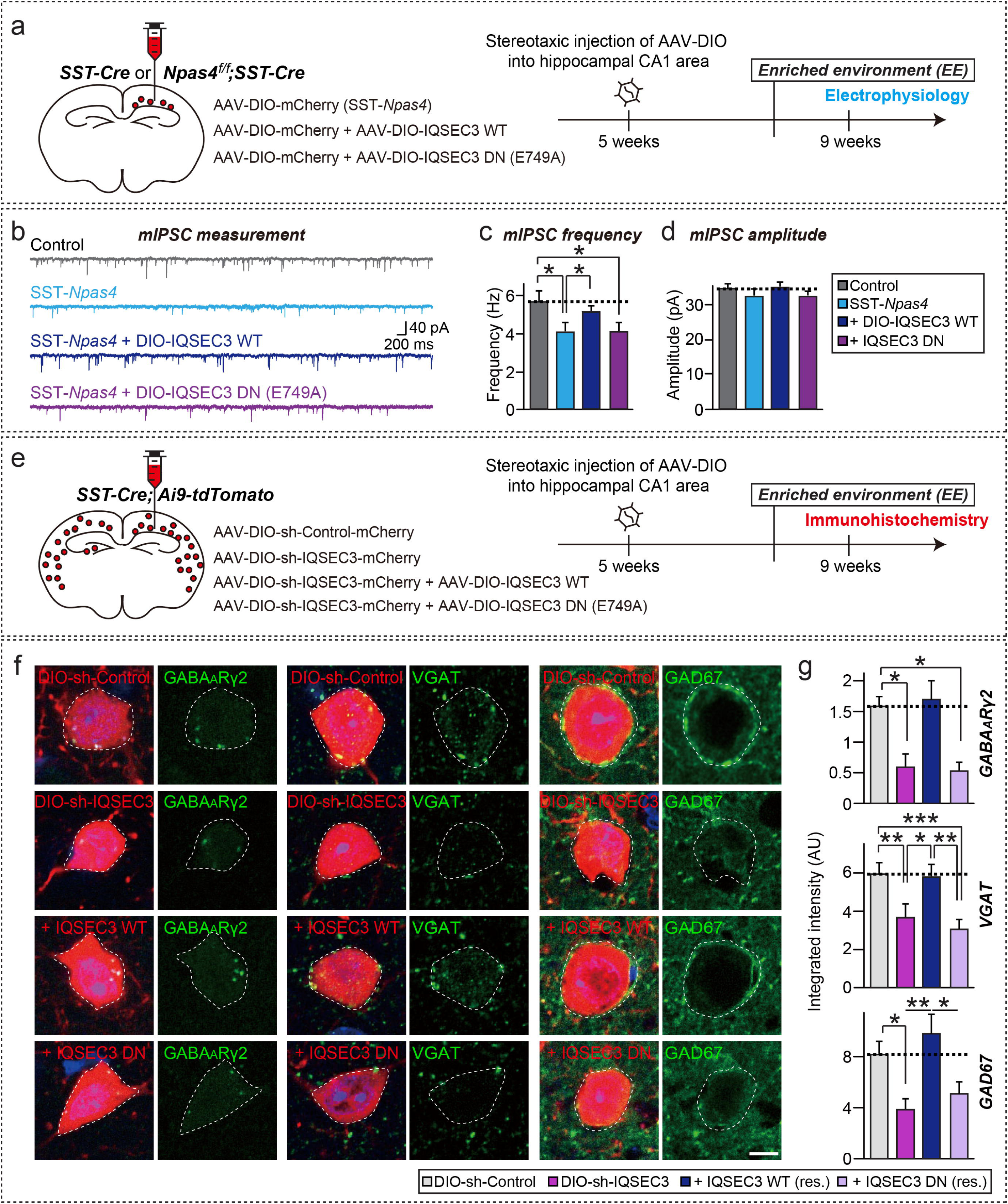
IQSEC3 is postsynaptically required for GABAergic synaptic transmission in somatostatin-expressing interneurons. **a** Experimental configuration for *ex vivo* electrophysiology experiments (**b–f**). **b–d** Representative mIPSC traces (**b**) and quantification of results (**c**, **d**) in neurons from SST-*Npas4*–KO mice exposed to EE conditions. Control, n = 29; SST-*Npas4,* n = 25; + DIO-IQSEC3 WT, n = 26; and +DIO-IQSEC3 DN, n = 34. Data shown are means ± SEMs (\**p* < 0.05; paired two-tailed t-test). See **Supplementary Table 2** for intrinsic electrophysiological properties of WT and SST-*Npas4–*KO neurons. **e** Experimental configuration for immunohistochemistry experiments (**h**, **i**). **f, g** Representative images (**f**) and quantification (**g**) of somatic GABA_A_Rγ2 (left), VGAT (middle), or GAD67 (right) puncta in the SO of neurons from SST-*Npas4*–KO mice exposed to EE conditions. Data shown are means ± SEMs (\**p* < 0.05; \*\**p* < 0.01, \*\*\**p* < 0.001; non-parametric ANOVA with Kruskal-Wallis test, followed by *post hoc* Dunn’s multiple comparison test; n = 4 mice for all conditions). Scale bar, 10 μm (applies to all images).

We then performed voltage-clamp recordings from SST^+^ (tdTomato^+^) neurons to assess predicted changes in GABAergic synaptic currents. Recordings from SST^+^ neurons of the hippocampal CA1 region revealed drastic reductions in the frequencies (but not amplitudes) of miniature inhibitory postsynaptic currents (mIPSCs) in SST-*Npas4–*KO mice (**Figure 4b–d**), in keeping with the loss of postsynaptic GABAγRs. Again, restricted introduction of IQSEC3 WT, but not IQSEC3 E749A, into SST-*Npas4–*KO mice rescued the deficits in GABAergic synaptic transmission (**Figure 4b–d**). The input resistance, resting membrane potentials, and resting membrane potentials of Npas4-deficient SST^+^ neurons were comparable to those of control neurons (**Supplementary Table 2**).

In addition, to determine whether IQSEC3 levels in SST^+^ neurons have an impact on the number of GABAergic synapses, we stereotactically injected AAV-sh-IQSEC3 into the hippocampal CA1 of SST-Cre; Ai9 mice (**Figure 4e**). Semi-quantitative immunohistochemical analyses showed that the overlap in the area of punctate immunoreactivity of the GABA_A_Rγ2 subunit (GABA_A_Rγ2) with the tdTomato^+^ somata of SST^+^ neurons in the SO layer of the hippocampus CA1 region was markedly reduced in SST-*Npas4–*KO mice compared with that in control mice (**Figure 4f, 4g**). Remarkably, transduction of Npas4-deficient SST^+^ neurons with double-floxed, inverted open reading frame (DIO), Cre-dependent AAVs expressing IQSEC3 WT, but not those expressing the ARF-GEF–inactive IQSEC3 E749A mutant completely reversed the decreased density of GABA_A_Rγ2 puncta in the SO (**Figure 4f, 4g**). Parallel analyses using VGAT or GAD67 antibodies yielded similar results (**Figure 4f, 4g**). These data suggest that IQSEC3 in SST^+^ CA1 hippocampal neurons postsynaptically regulates Npas4-mediated activity-dependent GABAergic synaptic transmission *in vivo*.

### IQSEC3 is required for Npas4-mediated GABAergic synaptic transmission in dendrites of hippocampal CA1 pyramidal neurons

Next, to determine if IQSEC3 is an Npas4 target in regulating somatic and/or dendritic inhibition of pyramidal neurons *in vivo*, we caused excision of Npas4 *in vivo* by injecting AAV-EGFP/Cre (Npas4 KO) into Npas4*^f/f^* mice; mice injected with AAV-EGFP/ΔCre served as controls. We then exposed these mice to EE for 4–5 days (**Figure 5a**), and recorded layer-specific, monosynaptically isolated, evoked inhibitory postsynaptic currents (eIPSCs) from AAV-infected CA1 pyramidal neurons (**Figure 5b–i**). Consistent with a previous report^8^, stimulation of axons in the SP layer produced marginally, but significantly, smaller eIPSCs in Npas4 KO neurons than in neighboring Npas4 WT neurons, whereas stimulation of axons in the stratum radiatum (SR) layer generated significantly larger eIPSCs in Npas4 KO neurons (**Figure 5b–e**). Strikingly, expression of IQSEC3 WT in Npas4 KO neurons completely reversed the increase in eIPSC amplitude induced by stimulation of axons in the SR layer, but did not affect the decrease in eIPSC amplitude induced by stimulation of axons in the SP layer, compared with uninfected Npas4 KO neurons (**Figure 5f, 5g**). Expression of IQSEC3 E749A in Npas4 KO neurons did not affect the amplitudes of eIPSCs induced by stimulation of axons in the SR or SP layer (**Figure 5h, 5i**). Paired-pulsed ratios (PPRs) measured following stimulation of axons in SR and SP layers of Npas4 WT and Npas4 KO mice housed under enriched-environment conditions and infected with AAVs expressing IQSEC3 WT or IQSEC3 E749A were not different among groups (**Supplementary Figure 4**), indicating that the Npas4-dependent changes in eIPSCs are not caused by altered presynaptic neurotransmitter release. These data indicate that Npas4 acts through IQSEC3 in a manner that depends on IQSEC3 ARF-GEF activity to mediate dendritic GABAergic synaptic inhibition *in vivo*.

**Fig. 5.**
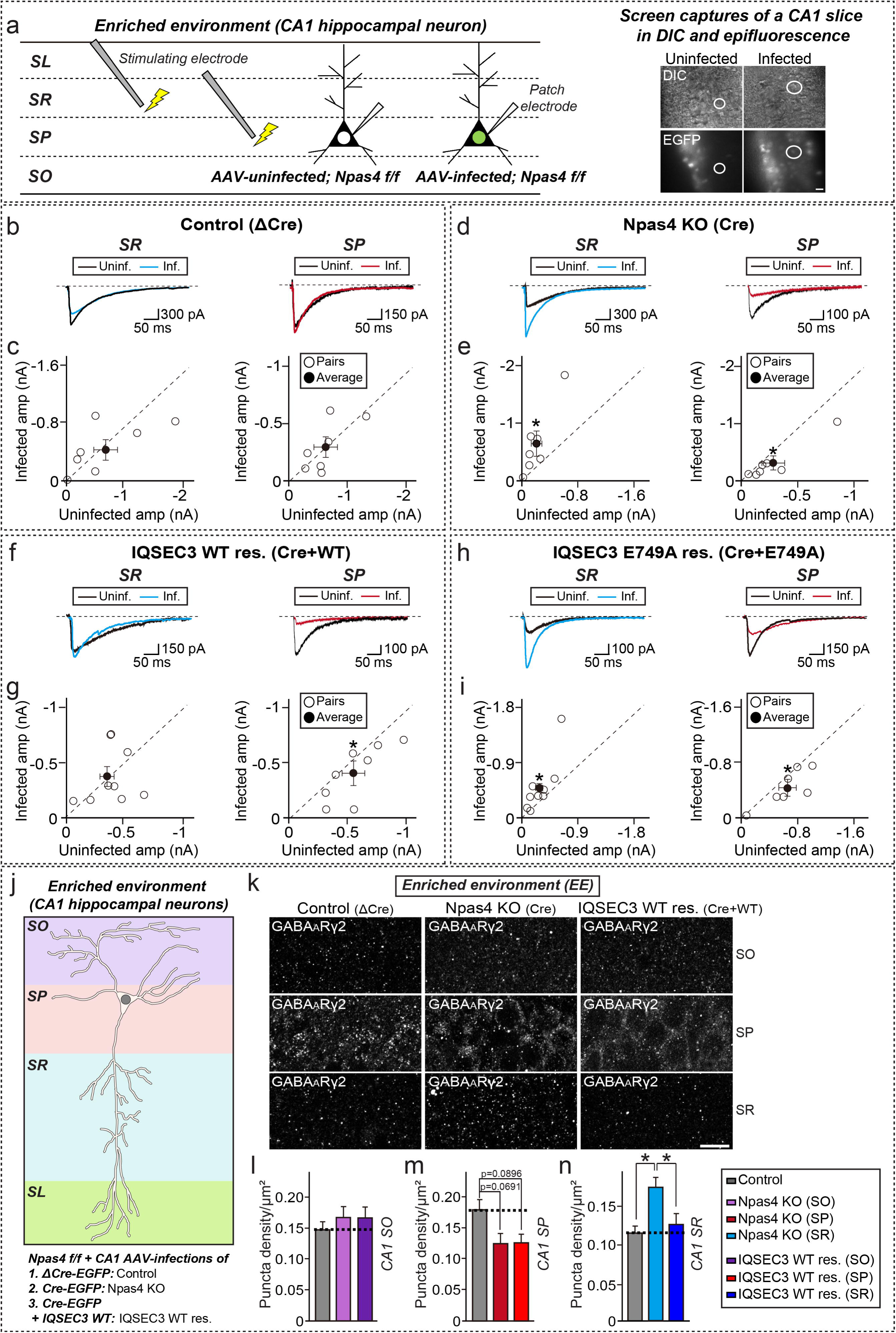
IQSEC3 regulates behaviorally induced, Npas4-mediated dendritic, but not somatic, inhibition. **a** Experimental configuration for *ex vivo* electrophysiology experiments. SL, stratum lacunosum; SR, stratum radiatum; SP, stratum pyramidale; and SO, stratum oriens. **b–i** eIPSC measurements in mice exposed to EE. Representative traces (**b**, **d**, **f**, **h**) of responses in neurons infected with the indicated AAVs (blue or red) to stimulation in the SR or SP layer, normalized pairwise to responses in uninfected neighboring neurons (black). Quantification results are presented in (**c**, **e**, **g**, **i**). Open circles represent Npas4 KO/Npas4 WT pairs; black circles indicate means ± SEM (SR [blue]: Control (ΔCre), n = 7 pairs; Npas4 KO (Cre), n = 7 pairs; WT rescue (Cre + IQSEC3 WT), n = 9 pairs; E749A rescue (Cre + IQSEC3 E749A), n = 10 pairs. SP [red]: Control (ΔCre), n = 7 pairs; Npas4 KO (Cre), n = 7 pairs; WT rescue (Cre + IQSEC3 WT), n = 8 pairs; and E749A rescue (Cre + IQSEC3 E749A), n = 7 pairs). Data shown are means ± SEMs (\**p* < 0.05; paired two-tailed t-test). See **Supplementary Table 3** for intrinsic electrophysiological properties of Npas4 WT and Npas4 KO neurons. **j** Experimental configuration for *in vivo* imaging experiments. SL, stratum lacunosum; SR, stratum radiatum; SP, stratum pyramidale; and SO, stratum oriens. **k–n** Representative images (**k**) and quantification of dendritic GABA_A_Rγ2 puncta in the SO (**l**) or SR (**m**), and somatic GABA_A_Rγ2 puncta in the SP (**n**) of neurons obtained from mice housed under EE conditions and infected with either AAV-ΔCre-EGFP (Control), AAV-Cre-EGFP (Npas4 KO), or AAV-Cre-EGFP plus AAV-IQSEC3 WT (IQSEC3 WT res.). Data are presented as means ± SEMs (\**p* < 0.05; non-parametric ANOVA with Kruskal-Wallis test, followed by *post hoc* Dunn’s multiple comparison test. n = 18**–**24 sections/3 mice for all conditions). Scale bar, 10 μm (applies to all images).

To corroborate the electrophysiological data obtained from CA1 Npas4 KO neurons (**Figure 5b–i**), we performed semi-quantitative analyses of endogenous GABA_A_Rγ2 puncta in the soma (SP layer), apical dendrites (SR layer), and basal dendrites (SO layer) of CA1 hippocampal neurons of Npas4*^f/f^* mice coinjected with AAVs encoding IQSEC3 WT and either EGFP-Cre (Npas4 KO) or EGFP-ΔCre (Control) (**Figure 5j**). Infected neurons were labeled by expressing AAV-EGFP-Cre or AAV-EGFP-ΔCre, whereas the differential effects of IQSEC3 expression in both somatic (SP layer) and dendritic (SR or SO layer) compartments on GABAergic synapse development were quantified by expressing AAV-IQSEC3 WT at moderate levels. The number of GABA_A_Rγ2 puncta in the SP layer of Npas4 KO neurons from mice exposed to enriched environment (EE) was significantly decreased compared with that in control neurons (**Figure 5k–n**), with a concomitant increase in the number of apical dendritic GABAARγ2 puncta in the SR (but not SO) layer of Npas4 KO neurons, consistent with a GABA_A_Rγ2 puncta in the SR (but not SO) layer of Npas4 KO neurons, consistent with a previous report^8^. Remarkably, IQSEC3 WT expression in Npas4 KO neurons completely abrogated the increase in the density of GABA_A_Rγ2 puncta in the SR only in mice exposed to EE conditions, without altering SP GABA_A_Rγ2 puncta density (**Figure 5k–n**), consistent with the electrophysiology data. These data suggest that IQSEC3 exerts differential Npas4 expression-dependent inhibition *in vivo*.

### IQSEC3 is required for controlling hippocampal CA1- and Npas4-dependent anxiety-like behavior

Lastly, we investigated the behavioral effects of IQSEC3 expression in Npas4 KO mice. Npas4 KO mice have been reported to exhibit diverse cognitive, emotional, and social behavioral deficits^10, 18, 36–38^. To link the hippocampal CA1 electrophysiological phenotypes in Npas4 KO mice with behavioral abnormalities, we performed a number of behavioral tasks, including elevated plus maze (EPM), Y-maze, open-field (OF), novel object-recognition (NOR), and forced swim (FS) tests using SST-*Npas4* KO and littermate control mice. SST-*Npas4* KO mice were additionally injected in the hippocampal CA1 region with AAV-DIO-mCherry (Npas4 cKO), AAV-DIO-IQSEC3 WT or AAV-DIO-IQSEC3 E749A (**Figure 6a**). SST-specific *Npas4*-KO mice exhibited decreased anxiety/exploration-related behavior (increased entries with similar time spent in open arms in the EPM test; **Figure 6b–d**), but normal depressive related behavior (as assessed by FS test; **Supplementary Figure 5**), normal locomotor activity (as assessed by the OF test; **Supplementary Figure 6**), normal working memory (as assessed by the Y-maze test; **Supplementary Figure 7**), and normal recognition memory (as assessed by the NOR test; **Supplementary Figure 8**) phenotypes that are slightly different from those of constitutive Npas4 KO mice^37, 38^. Strikingly, expression of IQSEC3 WT in CA1 SST neurons was sufficient to rescue the altered anxiety-related behavior (EPM test) observed in SST-specific *Npas4*-KO mice (**Figures 6b–d**). Intriguingly, expression of IQSEC3 E749A failed to normalize the increased anxiety-related behavior, suggesting that the ARF-GEF activity of IQSEC3 is required for mediating the ability to form a subset of Npas4- and CA1 SST-dependent mouse behaviors (**Figures 6b–d**). Expression of BDNF in CA1 SST neurons in the SST-specific *Npas4*-KO mice induced enormous hyperactivity and anxiety, consistent with previous reports^39, 40^, thus, we were unable to perform Y-maze, EPM, or FS tests on BDNF-expressing SST-specific *Npas4*-KO mice (**data not shown**). Taken together, together with previous studies^15, 18^, these data suggest that Npas4 employs distinct sets of downstream components in its roles in various types of cognitive tasks involving different brain regions.

**Fig. 6.**
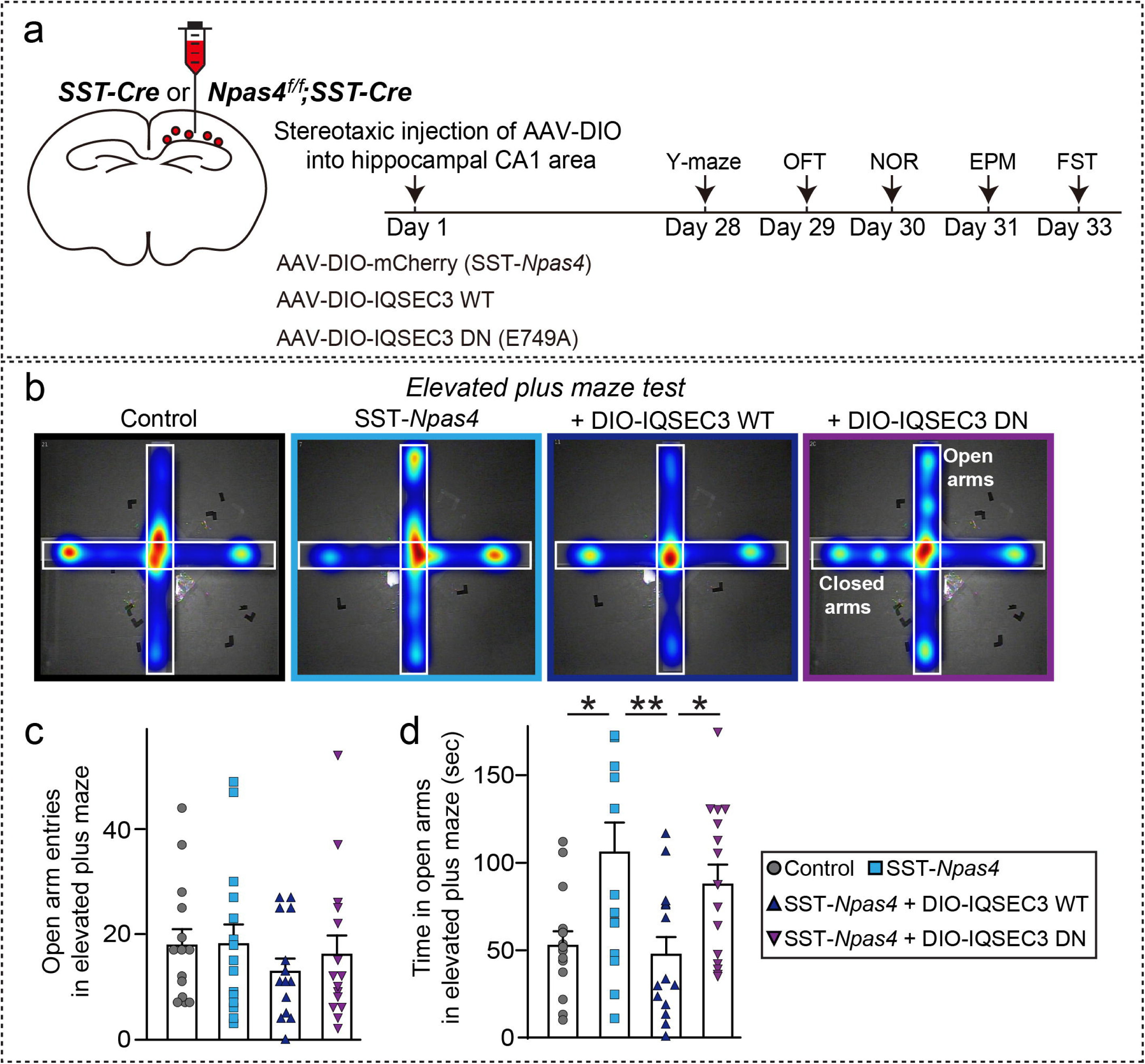
Expression of IQSEC3 WT, but not IQSEC3 E749A, in the hippocampal CA1 normalizes altered anxiety-like behavior observed in KO mice lacking Npas4 in somatostatin-positive neurons. **a** Schematic diagram of mouse behavioral analyses. The CA1 region of the hippocampus of ∼6-week-old SST-*Npas4* mice was bilaterally injected with AAVs expressing mCherry (SST-*Npas4* KO), or either of two IQSEC3 DIO-AAVs [AAV-DIO-IQSEC3 WT or AAV-DIO-IQSEC3 DN (E749A)]. WT littermate mice were used as controls. Injected mice were subjected to five behavioral tasks 4 weeks after the injections in the indicated order. Abbreviations: EPM, elevated plus-maze test; FS, forced swim test; NOR, novel object-recognition test; and OF, open-field test. **b–d** Analysis of anxiety/exploration-related behavior by EPM test in SST-*Npas4* mice injected with the indicated AAVs. Representative heat maps of the time spent in the open arms of the EPM (**b**) are presented. Red represents increased time spent, and blue represents minimal time spent during the test. Number of open arm entries (**c**) and time spent (**d**) in open arms were measured. Data are presented as means ± SEMs (Control, n = 14; SST-*Npas4*, n = 15; Rescue [+IQSEC3 WT], n = 14; and Rescue [+IQSEC3 E749A], n = 15; **\****p* < 0.05; \*\**p* < 0.01, one-way ANOVA with Bonferroni’s *post hoc* test).

## Discussion

In this study, we provide evidence from various experimental approaches to support the conclusion that IQSEC3 organizes activity-dependent GABAergic synapse development and controls hippocampal CA1-dependent anxiety-like behavior through intricate molecular mechanisms involving Npas4 and ARFs.

IQSEC3 is linked to Npas4-mediated post-transcriptional programs that orchestrate activity-dependent GABAergic synapse development. The activity-dependent regulation of IQSEC3 expression level prompted us to dissect how IQSEC3 is related to the previously elucidated mechanisms underlying activity-dependent GABAergic synapse development. A prime candidate was Npas4, a well-known transcription factor and putative upstream regulator of IQSEC3^8, 10^. Npas4 binds to the promoter region of *Iqsec3*, which contains Npas4 consensus binding sites; IQSEC3 protein, in turn, functions downstream of Npas4 in regulating GABAergic inhibition in both somatic and dendritic compartments of cultured hippocampal neurons (**Figure 1** and **Supplementary Figure 1**). IQSEC3 expression is specifically induced by manipulation of synaptic activity *in vivo* (**Figure 2** and **Supplementary Figure 3**), consistent with the idea that IQSEC3, whose corresponding gene is an Npas4 target, promotes activity-dependent GABAergic synapse development. This interpretation is further supported by the observations that upregulated IQSEC3 protein levels in SST^+^ neurons of the hippocampal CA1 SO layer were completely compromised in SST-specific *Npas4*-KO mice (**Figure 2**, 3 and **Supplementary Figure 3**). Although prior screens using GABAergic neuron-enriched cultures to profile activity-dependent transcriptomics and single-cell analyses in the mouse visual cortex failed to identify IQSEC3 as a stimulus-responsive gene^9, 41^, experimental conditions employed in these previous studies might have been unsuitable for identifying responsive genes, that are specifically regulated in hippocampal CA1 SST interneurons. Our results also suggest that context-dependent regulatory mechanisms may underlie Npas4-mediated transcriptional and post-transcriptional control of IQSEC3 levels *in vivo*^42^. Notably, Npas4 differentially dictates gene regulatory mechanisms tailored to the type of activity patterns along subcellular compartments of a neuron^43^.

IQSEC3 performs its inhibitory actions, distinct from other known Npas4 downstream components that are involved in regulating neural circuit-wide homeostasis (BDNF and Homer1a), neuroprotective signaling (Syt10), and synaptic plasticity during memory formation (Plk2) across various brain regions, neuron types and synapse types^9, 17, 18, 44^. Loss of Npas4 function results in reduced GABAergic synapse density in both perisomatic and dendritic regions of cultured hippocampal neurons, and blunts the inhibitory mode, concurrent with an abnormal distribution of GABAergic synapses at both soma and apical dendrites of CA1 pyramidal neurons^7^. BDNF is specifically expressed in excitatory ‘cortical’ neurons and serves to specifically promote perisomatic inhibition of them, whereas Plk2 is strongly expressed in excitatory ‘hippocampal’ CA3 pyramidal neurons that regulate thorny excrescence structures at mossy fiber-CA3 synapses^9, 18^. Anatomical analyses in the current study revealed that IQSEC3 KD abolished the Npas4-induced increase in inhibitory synaptic puncta density at both perisomatic and apical dendritic regions of hippocampal cultured neurons, whereas IQSEC3 overexpression in SST-specific *Npas4*-KO mice resulted in selective restoration of altered inhibitory synaptic transmission and structure in apical dendritic regions in CA1 hippocampal pyramidal neurons (**Figures 4**, **5**, and **Supplementary Figure 1**). This mechanism of IQSEC3 action is in stark contrast to the specific mode of action of BDNF and neuroligin-2 at perisomatic regions of hippocampal neurons^7, 8, 45^. These observations might be integrated under a scenario in which altered inhibitory synapse development in both somatic and dendritic compartments of IQSEC3-deficient neurons involves at least two distinct pathways *in vivo*: an Npas4-dependent IQSEC3 pathway that mediates dendritic inhibition (**Figure 7b**), and an Npas4-independent IQSEC3 pathway that mediates somatic and/or dendritic inhibition (**Figure 7a**). Consistent with this speculation, activity-dependent upregulation of IQSEC3 was notable in SST interneurons (but not excitatory neurons) of the hippocampal CA1 region (**Figure 2**), suggesting that one plausible role of IQSEC3 in SST neurons is to increase inhibitory inputs onto them enabling Npas4 to manage optimal responses and enhance feedback inhibition of principal neurons within local neural circuits in a context-dependent fashion (**Figure 7**). Given that IQSEC3 is expressed in various types of interneurons (PV^+^, SST^+^, or CCK^+^) in mouse hippocampal regions (**data not shown**), it is likely that IQSEC3 differentially shapes specific circuit features that modulate homeostatic plasticity at distinct principal neuron-interneuron synapses. Indeed, SST^+^ interneurons also provide inhibitory input to PV^+^ interneurons, which preferentially innervate the perisomatic region and axon initial segment of pyramidal cells, thereby controlling their spike output^45^. Thus, it is possible that increased inhibition of PV^+^ cells by disinhibited SST+ neurons might also contribute to disinhibition of pyramidal cells in SST-*Npas4* KO mice. In support of this conjecture, increased eIPSC amplitudes were observed in Npas4-deficient pyramidal cells of the SP layer (**Figure 5**). Future studies in which Npas4 is conditionally deleted in PV^+^ interneurons are warranted to clarify this issue.

**Fig. 7.**
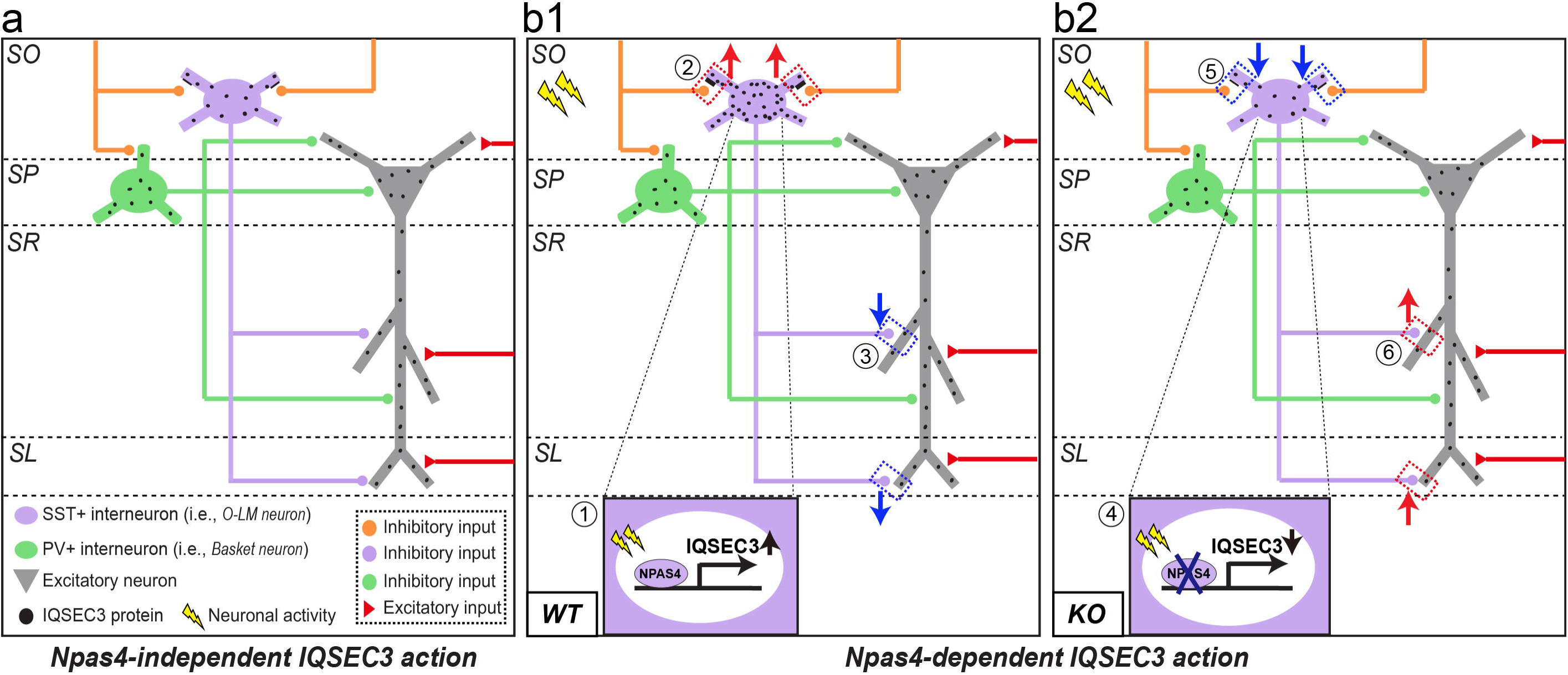
Molecular model of IQSEC3 action in regulating Npas4 activity-dependent neural circuit-wide homeostasis. **a** Npas4-independent IQSEC3 action in controlling activity-dependent GABAergic synapse development across universal synapse types. **b** Npas4-dependent IQSEC3 action in controlling activity-dependent GABAergic synapse development in specific synapse types. Neuronal activity-induced transcriptional programs are orchestrated by coordinated actions of various Npas4 downstream genes in distinct neuron types to maintain proper network activity. In WT mice (**b1**), a specific pattern of neuronal activity (①) specifically upregulates IQSEC3 expression in SST interneurons, possibly inducing increased inhibitory input onto SST interneurons (②), and act in conjunction with decreased inhibitory drive onto excitatory neurons to maintain circuit activity (③). This occurs despite widespread upregulation of Npas4 expression in both excitatory and inhibitory neurons, which may activate other Npas4 target genes in both excitatory and inhibitory neurons to maintain circuit activity. In contrast, neuronal activity fails to induce IQSEC3 upregulation in SST interneurons of Npas4 KO mice (**b2)** (④), possibly resulting in reduced recruitment of inhibitory inputs onto SST interneurons (⑤), and collectively contributing to increased inhibitory drive onto a specific dendritic domain in excitatory neurons (⑥), namely the dendritic domain in the SR layer, but not the SL or SO layer, of the hippocampal CA1 region).

Expression of IQSEC3 in CA1 SST interneurons completely normalized the altered anxiety-like behavior of SST-specific *Npas4*-KO mice (**Figure 6**), suggesting that a distinct set of factors downstream of Npas4 is required for the modulation of a spectrum of hippocampus-dependent cognitive and behavioral tasks (e.g., IQSEC3 for CA1-dependent anxiety-like behavior, and BDNF and Plk2 for CA3-dependent contextual memory formation). More importantly, our results suggest that the altered anxiety-like behavioral phenotype of SST-*Npas4* KO mice reflects differential GABAergic inhibition of pyramidal cells by the context-dependent, balanced activity of SST+ interneurons and PV+ interneurons. Conditional transgenic mice in which IQSEC3 is selectively deleted in specific interneurons should be further analyzed to address how various neural circuits involving different cell types are altered in the context of the GABAergic synaptic functions of IQSEC3 proposed in the current study. This further study is especially meaningful given the implications of SST^+^ GABAergic neurons in modulating anxiolytic brain states and major depressive disorders through the control of inhibition at specific synapse types formed in discrete neural circuits^47,48,49,50^. Other intriguing questions include whether hyper-excitability of SST^+^ neurons leads to increased expression and release of SST peptides that might contribute to the behavioral phenotype of SST-*Npas4* KO mice, and how ARF signaling pathways are differentially involved in cell type-specific manner.

In conclusion, our observations underscore the significance of biochemical complexes involving IQSEC3, Npas4, ARFs, and other components in organizing activity-dependent GABAergic synapse development, and in shaping specific neural circuit architectures involved in a specific behavior.

## Supporting information

Manuscript

## Acknowledgements.

We are grateful to Drs. Yingxi Lin (MIT, USA) and Hiroyuki Sakagami (Kitasato University, Japan) for kind gifts of reagents. This study was supported by grants from Future Planning (2019R1A2B5B02069324 to J.K.; 2019R1A2C1086048to J.W.U), the Brain the National Research Foundation of Korea (NRF) funded by the Ministry of Science and Future Planning (2019R1A2B5B02069324 to J.K.; 2019R1A2C1086048to J.W.U), the Brain Research Program through the NRF funded by the Ministry of Science, ICT & Future Planning (2017M3C7A1023470 to J.K. and 2017M3C7A1023471 to E.C.). The authors declare no competing financial interests.

## Author contributions

J.W.U. and J.K. conceived and directed the project; S.K., D.P., Jinhu K., D.L., D.K., H.K., S.H. and J.J. performed the experiments; S.K., D.P., Jinhu K., D.L., Jaehoon K., E.C., J.W.U. and J.K. analyzed the data; and E.C., J.W.U. and J.K. wrote the manuscript. All authors read and commented on the manuscript.

## Methods

### Cell culture

HEK293T cells were cultured in Dulbecco’s Modified Eagle’s Medium (DMEM; WELGENE) supplemented with 10% fetal bovine serum (FBS; Tissue Culture Biologicals) and 1% penicillin-streptomycin (Thermo Fisher) at 37°C in a humidified 5% CO_2_ atmosphere. Cultured primary hippocampal neurons were prepared from embryonic day 18 (E18) Sprague-Dawley rat brains (KOATECK). Neurons were seeded on 25-mm poly-L-lysine (1 mg/ml)– coated coverslips and cultured in Neurobasal media (Gibco) containing penicillin-streptomycin and 0.5 mM GlutaMax (Thermo Fisher) supplemented with 2% B-27 (Thermo Fisher) and 0.5% FBS (Hyclone). All procedures were conducted according to the guidelines and protocols for rodent experimentation approved by the Institutional Animal Care and Use Committee of DGIST.

### Animals

All C57BL/6N mice were maintained and handled in accordance with protocols approved by the Institutional Animal Care and Use Committee of DGIST under standard, temperature-controlled laboratory conditions, or in an enriched environment with free access to colored tunnels, mazes, climbing materials, and running wheels. Mice were kept on a 12:12 light/dark cycle (lights on at 6:00 am), and received water and food *ad libitum.* Npas4 KO (Npas4^*f/f*^) mice were previously described7. Ai9 reporter mice (007909, Jackson Research Laboratories) and Sst-ires-Cre mice (013044, Jackson Research Laboratories) were gifts of Dr. Eunjoon Kim (KAIST, Korea). All experimental procedures were performed on male mice. Pregnant rats purchased from Daehan Biolink were used *in vitro* for dissociated cultures of cortical or hippocampal neurons.

### Construction of expression vectors

*1.* IQSEC3. AAVs encoding full-length rat *Iqsec3* WT and the E749A point mutant were generated by amplification of the full-length region by PCR and subsequent subcloning into the pAAV-T2A-tdTomato vector (a gift from Hailan Hu^51^) at *Xba*I and *BamH*I sites. DIO-rat *Iqsec*3 WT and E749A fragments were PCR amplified and subcloned into the pAAV-hSynI-DIO-mCherry vector at *Nhe*I sites. The shRNA AAV against mouse *Iqsec3* (GenBank accession number: NM_207617.1) was constructed by annealing, phosphorylating, and cloning oligonucleotides targeting mouse *Iqsec3* (5’-GAA CTG GTG GTA GGC ATC TAT GAG A-3’) into the *BamH*I and *EcoR*I sites of the pAAV-U6-GFP vector (Cell BioLabs, Inc.) or *Avr*II and *EcoR*I sites of the pAAV-Ef1α-DSE-mCherry-PSE vector (Addgene). *2*. Others. The shRNA lentiviral expression vector against *Bdnf* was constructed by annealing, phosphorylating, and cloning oligonucleotides targeting rat *Bdnf* (Genbank accession number: AY559250; 5’-ACA TCA GAC CAT CGG CCA CGA CATT A-3’) into the *Xho*I and *Xba*I sites of the L-315 vector. AAV encoding mouse *Bdnf* (Genbank accession number: AY231132) was generated by amplification of the full-length region by PCR and subsequent subcloning into the pAAV-hSynI-DIO-mCherry vector at *Nhe*I sites. *3*. Previously published reagents. The following constructs were as previously described: pcDNA3.1-Myc-Npas4 [a gift from Yingxi Lin]^7^; L-315 rat sh-IQSEC3, pcDNA3.1-Myc-IQSEC3 (shRNA-resistant) WT and E749A^24^; pAAV2/9-Cre-IRES-EGFP and pAAV2/9-ΔCre-IRES-EGFP^52^; and pFSW-Cre and pFSW-ΔCre^53^.

### Antibodies

The following commercially available antibodies were used: goat polyclonal anti-EGFP (Rockland), mouse monoclonal anti-Npas4 (clone N408/79; Neuromab), rabbit polyclonal anti-cholecystokinin-8 (Immunostar), mouse monoclonal anti-GAD67 (clone 1G10.2; Millipore), guinea pig polyclonal anti-VGLUT1 (Millipore), rabbit polyclonal anti-VGAT (Millipore), mouse monoclonal anti-gephyrin (clone 3B11; Synaptic Systems), rabbit polyclonal anti-GABA_A_ receptor γ2 (Synaptic Systems), mouse monoclonal anti-PSD-95 (clone 7E3-1B8; Thermo Fischer Scientific), mouse monoclonal anti-β-actin (clone C4; Santa Cruz Biotechnology), mouse monoclonal anti-parvalbumin (clone PARV-19; Swant), and mouse monoclonal anti-somatostatin (clone YC7; Millipore). The following antibodies were previously described: rabbit polyclonal anti-IQSEC3 [JK079]^24^; guinea pig polyclonal anti-IQSEC3/SynArfGEF [a gift from Hiroyuki Sakagami]^27^; and rabbit polyclonal anti-Npas4 [a gift from Michael Greenberg]^8^.

### *Iqsec3* promoter-luciferase reporter analysis

A DNA fragment bracketing the proximal promoter and transcription start site of the *Iqsec3* gene (–280 to +36) was PCR-amplified from mouse genomic DNA and subcloned into the pGL3-Basic vector (Promega). Full-length mouse Npas4 cDNA was subcloned in pcDNA3 (Gibco-Invitrogen). For luciferase assays, 5 × 10^4^ HEK293T cells or cultured hippocampal neurons in 24-well plates were transfected with an Npas4 expression vector and reporter plasmid using iN-fect (iNtRON Biotechnology; for HEK293T cells) or CalPhos (Clontech; for cultured hippocampal neurons). Cells were harvested after 48 hours and analyzed for luciferase activity as described by the manufacturer (Promega).

### Electrophoretic mobility shift assay

Full-length mouse Npas4 cDNA was subcloned into the pFASTBAC1 vector containing an N-terminal FLAG-tag, and baculovirus was generated according to the manufacturer’s instructions (Thermo Fisher). After baculovirus infection into Sf9 insect cells, Npas4 protein was affinity purified on M2 agarose (Sigma) as described previously^54^. An oligonucleotide containing the consensus sequence for Npas4 binding (5’-CGG GAA GCA GGG TTC CTC ACG AAG CCC AGA GGT GGG AGG TGG-3’) and its point mutant (5’-CGG GAA GCA GGG TTC CT<u>A AAA</u> AAG CCC AGA GGT GGG AGG TGG-3’; mutated nucleotides are underlined) within the *Iqsec3* proximal promoter region were end-labeled using polynucleotide kinase and [γ-^32^P]ATP and then annealed to complementary oligonucleotides. Reactions containing purified Npas4 protein in 19 µl of reaction buffer (10 mM Tris-Cl pH 7.5, 5 mM MgCl_2_, 1 mM EDTA, 50 mM K-glutamate, 1 mM DTT and 5% glycerol) supplemented with 25 ng/μl poly (dI-dC) were pre-incubated on ice for 15 minutes. After addition of 1 µ l of ^32^P-labeled probe containing approximately 3 ng of oligonucleotide duplex, reactions were incubated on ice for 30 minutes. Protein in samples were resolved by electrophoresis at 4°C on 6% polyacrylamide gels in 0.25x TBE buffer and subjected to autoradiography.

### Chromatin immunoprecipitation assays

Cultured cortical neurons at DIV14 were treated with 55 mM KCl for 6 hours, after which chromatin immunoprecipitation assays were performed according the manufacturer’s instructions (Millipore). Detailed information about primers used for quantitative PCR is provided in **Supplementary Table 1**.

### Chemicals

Amino-5-phosphonopentanoic acid (D-APV; Cat. No. 0106), 2,3-dihydroxy-6-nitro-7-sulfamoyl-benzo[f] quinoxaline-2,3-dione (NBQX; Cat. No. 0373), 6,7-dinitroquinoxaline-2,3(1H,4H)-dione (DNQX; Cat. No. 0189), tetrodotoxin (TTX; Cat. No. 1078), and QX-314 (CNQX; Cat. No. C239), picrotoxin (Cat. No. P1675), 2,2,2-tribomoethyl alcohol (Cat. No. T4840-2), and Tert-amylalcohol (Cat. No. 24048-6), and kainic acid (Cat. No. K0250) were purchased from Sigma. Diazepam (Cat. No. 117) was purchased from Daewon Pharm.

### Neuron culture, transfections, imaging, and quantitation

Cultured hippocampal neurons were prepared from E18 rat brains, as previously described^55, 56^, cultured on coverslips coated with poly-D-lysine (Sigma), and grown in Neurobasal medium supplemented with B-27 (Thermo Fisher), 0.5% FBS (WELGENE), 0.5 mM GlutaMAX (Thermo Fisher), and sodium pyruvate (Thermo Fisher). For overexpression of IQSEC3 in cultured neurons, hippocampal neurons were transfected with pCAGGS-FLAG-IQSEC3 or its various derivatives, as indicated in the individual figures, or with EGFP (Control) using a CalPhos Transfection Kit (Takara) at DIV10 and immunostained at DIV14. For KD of IQSEC3 in cultured neurons, hippocampal neurons were transfected with L-315 alone (Control), L-315 rat sh-IQSEC3 (IQSEC3 KD), or cotransfected with IQSEC3 KD and shRNA-resistant myc-IQSEC3 using a CalPhos Transfection Kit (Takara) at DIV8 and immunostained at DIV14. For immunocytochemistry, cultured neurons were fixed with 4% paraformaldehyde/4% sucrose, permeabilized with 0.2% Triton X-100 in phosphate buffered saline (PBS), immunostained with the indicated primary antibodies, and detected with the indicated Cy3- and fluorescein isothiocyanate (FITC)-conjugated secondary antibodies (Jackson ImmunoResearch). Images were acquired using a confocal microscope (LSM780, Carl Zeiss) with a 63 x objective lens; all image setting were kept constant. Z-stack images were converted to maximal intensity projection and analyzed to obtain the size, intensity, and density of puncta immunoreactivities derived from marker proteins. Quantification was performed in a blinded manner using MetaMorph software (Molecular Devices).

### AAV production, stereotactic surgery and virus injection

Recombinant AAVs were packaged with pHelper and AAV1.0 (serotype 2/9) capsids for high efficiency. HEK293T cells were cotransfected with pHelper and pAAV1.0, together with pAAV-U6-EGFP alone (Control), pAAV-U6-shNpas4 (Npas4 KD), pAAV-U6-sh-IQSEC3 (IQSEC3 KD), pAAV-DIO-IQSEC3 WT, pAAV-DIO-IQSEC3 E749A, pAAV-IQSEC3 WT or pAAV-IQSEC3 E749A. Cells were harvested 72–108 hours post transfection; after adding 0.5 M EDTA to the media, cells were washed three times with PBS, and collected by centrifugation. Cells were then resuspended in PBS and lysed by subjecting them to four freeze-thaw cycles in an ethanol/dry ice bath (7 minutes each) and 37 °C water bath (5 minutes). Lysates were centrifuged and supernatants were collected and incubated with a solution containing 40% poly(ethylene glycol) (Sigma) and resuspended in HEPES buffer (20 mM HEPES pH 8.0, 115 mM NaCl, 1.2 mM CaCl_2_, 1.2 mM MgCl_2_, 2.4 mM KH_2_PO_4_), mixed with chloroform, and centrifuged at 400 rcf for 10 minutes. The supernatant was collected and concentrated using Amicon Ultra Centrifugal Filters (0.5 ml, 3K MWCO; Millipore). Viruses were assessed for infectious titer by RT-PCR, and used for infections at 1 × 10^10^–10^11^ infectious units/μl. For stereotactic delivery of recombinant AAVs, 6–8-week-old C57BL/6N mice were anesthetized by inhalation of isoflurane (3–4%) or intraperitoneal injection of 2% Avertin solution (2,2,2-tribromoethyl alcohol dissolved in Tert-amylalcohol (Sigma)) dissolved in saline, and secured in the stereotactic apparatus. Viral solutions were injected with a Hamilton syringe using a Nanoliter 2010 Injector (World Precision Instruments) at a flow rate of 100 nl/min (injected volume, 0.6 μl). The coordinates used for stereotactic injections into mice are as follows: hippocampal DG injections (anterior-posterior [AP] −2.2 mm, medial-lateral [ML] ± 1.3 mm, and dorso-ventral [DV] 2.35 mm) and hippocampal CA1 injections (AP −2.2 mm, ML ± 1.3 mm, and DV 1.9 mm). Each injected mouse was returned to its home cage and used for scoring seizure-like behaviors, immunohistochemical analyses, electrophysiological recordings or behavioral analyses after 2–4 weeks.

### qRT-PCR

Cultured rat cortical neurons were infected with recombinant lentiviruses at DIV3 and harvested at DIV10 for qRT-PCR using SYBR green qPCR master mix (Takara). Total RNA was extracted from rat cortical neurons using the TRIzol reagent (Invitrogen) according to the manufacturer’s protocol. Briefly, one well of a 12-well plate of cultured neurons was harvested and incubated with 500 μl of TRIzol reagent at room temperature for 5 minutes. After phenol-chloroform extraction, RNA in the upper aqueous phase was precipitated. cDNA was synthesized from 500 ng of RNA by reverse transcription using a ReverTra Ace-α kit (Toyobo). Quantitative PCR was performed on a CFX96 Touch Real-Time PCR system (BioRad) using 1 μl of cDNA. The ubiquitously expressed β-actin was used as an endogenous control. The sequences of the primer pairs used are as follows: rat I*qsec3*, 5’-GGA GCA GAT TCG GAT AGA ATG G-3’ (forward) and 5’–GGG TGA TCC TTG CTT TGA CT-3’ (reverse); rat *Npas4*, 5’-GAG GCT GGA CAT GGA TTT ACT–3’ (forward) and 5’-GGG TGA TCC TTG CTT TGA CT-3’ (reverse); rat *Bdnf*, 5’-CTG AGC GTG TGT GAC AGT ATT A-3’ (forward) and 5’-CTT TGG ATA CCG GGA CTT TCT C-3’; mouse *Npas4*, 5’-CCT CCA AAG AGC TGG ACT TC-3’ (forward) and 5’-ATC CTT GCT CAG GTC TGC TT-3’ (reverse); mouse *Iqsec3*, 5’-ATC ACC ACC AAC ATC ACC AC–3’ (forward) and 5’-CTT GAC AAT CTG CTG GCA CT-3’ (reverse); and mouse *Bdnf*, 5’-CTG AGC GTG TGT GAC AGT ATT A–3’ (forward) and 5’-CTT TGG ATA CCG GGA CTT TCT C-3’ (reverse)

### Activity alteration protocols

*<u>1. Environmental enrichment</u>.* Male Npas4*^f/f^* mice (P28–P30) injected with the indicated AAVs into the CA1 region were kept for 4 weeks and then exposed to an enriched environment for 4–5 days before functional analyses. The enriched environment consisted of a large cage containing a running wheel, hut, tunnel and several other novel objects, as previously described^57^. *<u>2. Induced seizure protocol</u>*. Seizures were induced by intraperitoneal administration of 11–12-week-old male mice with KA (15 mg/kg); saline injection was used as a control. Mice were decapitated at the indicated times (3 or 6 hours) after injection, and the injected mouse brains were prepared immediately for further analyses.

### Fluorescent *in situ* hybridization (RNAscope assay)

Frozen sections (14-µm thick) were cut coronally through the hippocampal formation and thaw-mounted onto Superfrost Plus microscope slides (Advanced Cell Diagnostics). Sections were fixed in 4% formaldehyde for 10 minutes, dehydrated in increasing concentrations of ethanol for 5 minutes, air-dried, and then pretreated with protease for 10 minutes at room temperature. For RNA detection, sections were incubated in different amplifier solutions in a HybEZ hybridization oven (Advanced Cell Diagnostics) at 40°C. Synthetic oligonucleotides complementary to the sequence corresponding to nucleotide residues 697–1045 of NM-001033354.3 (Mm–lqsec3–tv1), 1908–3117 of NM-001033354.3 (Mm-lqsec3–C2), 793-1795 of NM_153553.4 (Mm-Npas4-C2), and 18-407 of NM_009215.1 (Mm-SST-C3) (Advanced Cell Diagnostics) were used as probes. The labeled probes were conjugated to Alexa Fluor 488, Altto 550 or Altto 647, after which labeled probe mixtures were hybridized by incubating with slide-mounted sections for 2 hours at 40 °C. Nonspecifically hybridized probes were removed by washing the sections three times for 2 minutes each with 1X wash buffer at room temperature, followed by incubation with Amplifier 1-FL for 30 minutes, Amplifier 2–FL for 15 minutes, Amplifier 3–FL for 30 minutes, and Amplifier 4 Alt B–FL for 15 minutes at 40°C. Each amplifier was removed by washing with 1X wash buffer for 2 minutes at room temperature. The slides were imaged with an LSM780 microscope (Zeiss) and analyzed using Image J (NIH).

### Immunohistochemistry

Three-month-old mice were anaesthetized and immediately perfused, first with PBS for 3 minutes, and then with 4% paraformaldehyde for 5 minutes. Brains were dissected out, fixed in 4% paraformaldehyde overnight, and then incubated with 30% sucrose (in PBS) overnight, and sliced into 30-μm-thick coronal sections using a vibratome (Model VT1200S; Leica Biosystems) or a cryotome (Model CM-3050-S; Leica Biosystems). Sections were permeabilized by incubating with 1% Triton X-100 in PBS containing 5% bovine serum albumen and 5% horse serum for 30 minutes. For immunostaining, sections were incubated for 8–12 hours at 4 °C with primary antibodies diluted in the same blocking solution. The following primary antibodies were used: anti-Npas4 (1:100), anti-IQSEC3 (JK079; 2 μg/ml), anti-GAD67 (1:100), anti-PV (1:500), anti-CCK (1:100), and anti-SST (1:50). Sections were washed three times in PBS and incubated with appropriate Cy3- or FITC-conjugated secondary antibodies (Jackson ImmunoResearch) for 2 hours at room temperature. After three washes with PBS, sections were mounted onto glass slides (Superfrost Plus; Fisher Scientific) with Vectashield mounting medium (H-1200; Vector Laboratories).

### *Ex vivo* electrophysiology

*<u>1. SST interneuron recordings.</u>* Whole-cell voltage-clamp recordings were obtained from acute brain slices. Brain slices were transferred to a recording chamber and perfused with a bath solution of aerated (O_2_ 95%/CO_2_ 5% mixed gas) artificial cerebrospinal fluid (aCSF) consisting of 124 mM NaCl, 3.3 mM KCl, 1.3 mM NaH_2_PO_4_, 26 mM NaHCO_3_, 11 mM D-glucose, 2.5 mM CaCl_2_, and 1.5 mM MgCl_2_ at 30°C. Patch pipettes (open pipette resistance, 3–5_MΩ) were filled with an internal solution consisting of 145 mM CsCl, 5 mM NaCl, 10 mM HEPES, 10 mM EGTA, 4 mM Mg-ATP, and 0.3 mM Na-GTP. Whole-cell recordings of mIPSCs were performed on CA1 *stratum oriens* SST^+^ interneurons, voltage clamped at −70 mV, and currents were pharmacologically isolated by bath application of 50μM D-APV (Tocris), 10 μM CNQX (Sigma), and 1 μM TTX (Tocris). Electrophysiological data were acquired using pCLAMP software and a MultiClamp 700B (Axon Instruments), and were digitized using an Axon DigiData 1550B data acquisition board (Axon Instruments). Data were sampled at 10 kHz and filtered at 4 kHz. Experiments were discarded if the holding current was greater than −250 pA, the mean series resistance of pairs was greater than 30 ΩM, or the series resistance differed by more than 20% between the two recordings. *<u>2. Pyramidal neuron recordings</u>.* Whole-cell voltage-clamp recordings were obtained from acute brain slices. Brain slices were transferred to a recording chamber and perfused with a bath solution of aerated (O_2_ 95%/CO_2_ 5% mixed gas) artificial cerebrospinal fluid (aCSF) consisting of 124 mM NaCl, 3 mM KCl, 1.3 mM MgSO_4_, 1.25 mM NaH_2_PO_4_, 26 mM NaHCO_3_, 2.4 mM CaCl_2_-2H_2_O, and 10 mM glucose at 32°C. Patch pipettes (open pipette resistance, 3–5 MΩ) were filled with an internal solution consisting of 130 mM CsCl, 2 mM NgCl_2_, 10 mM HEPES, 5 mM Mg-ATP, 0.5 mM Na-GTP and 0.1 mM EGTA. For experiments in which eIPSCs were recorded, currents were pharmacologically isolated by bath application of 50 µM D-APV (Tocris) and 50 µM CNQX (Sigma), and inclusion of 5 mM QX-314 (Sigma) in the internal solution. Responses in hippocampal CA1 pyramidal neurons in the pyramidal layer evoked by stimulation of neurons in the SR layer were measured from a holding potential of −70 mV. Extracellular stimulation within specific layers of the hippocampus was achieved by current injection through a stimulating electrode placed in the relevant layer within 100–200 μm laterally of the patched pair. The stimulus strength used was the minimum required to generate an eIPSC in both Npas4 WT and Npas4 KO neurons. Electrophysiological data were acquired using pCLAMP software and a MultiClamp 700B (Axon Instruments), and were digitized using an Axon DigiData 1550B data acquisition board (Axon Instruments). Data were sampled at 10 kHz and filtered at 4 kHz. Experiments were discarded if the holding current was greater than −500 pA, the mean series resistance of pairs was greater than 30 MΩ, or if the series resistance differed by more than 30% between the two recordings. In experiments in which direct comparisons were made between two neurons, all data were normalized to the peak amplitude of Npas4 WT neurons. The amplitude of eIPSCs was calculated by averaging the normalized peak amplitude. Paired-pulse ratios were calculated by recording the amplitudes of paired eIPSC at stimulation intervals of 25, 50, 100 and 200 ms.

### Mouse behavioral tests

Male *Npas4*^f/f^ mice (9–11 weeks old) crossed with the SST-*ires*-Cre driver line were injected with the indicated AAVs were used for all behavioral tests. Tests were performed in the following order: Y-maze, open-field, novel object-recognition, elevated-plus maze, and forced swim test. Mice were excluded for quantitative analyses if one or both of the injections were off-target, as demonstrated by *post hoc* immunostaining after behavioral analyses. *<u>1. Y-maze test</u>.* A Y-shaped white acrylic maze with three 40-cm–long arms at a 120° angle from each other was used. Mice were introduced into the center of the maze and allowed to explore freely for 8 minutes. An entry was counted when all four limbs of a mouse were within the arm. The movement of mice was recorded by a top-view infrared camera, and analyzed using EthoVision XT 10 software (Noldus). *<u>2. Open-field test.</u>* Mice were placed into a white acrylic open-field box (40 × 40 × 40 cm), and allowed to freely explore the environment for 60 minutes in the dark (0 lux). The traveled distance moved and time spent in the center zone of freely moving mice were recorded by a top-view infrared camera, and analyzed using EthoVision XT 10 software (Noldus). *<u>3. Novel object-recognition test</u>*. An open field chamber was used in this test. Mice were habituated to the chamber for 10 minutes. For training sessions, two identical objects were placed in the center of the chamber at regular intervals, and mice were allowed to explore the objects for 10 minutes. After the training session, mice were returned to their home cage for 6 hours. For novel object-recognition tests, one of the two objects was exchanged for a new object, placed in the same position of the chamber. Mice were returned to the chamber and allowed to explore freely for 10 minutes. The movement of mice was recorded by infrared camera, and the number and duration of contacts were analyzed using EthoVision XT 10 (Noldus). *<u>4. Elevated plus-maze test.</u>* The elevated plus-maze is a plus-shaped (+) white acrylic maze with two open arms (30 × 5 × 0.5 cm) and two closed arms (30 × 5 × 30 cm) positioned at a height of 75 cm from the floor. Light conditions around open and closed arms were ∼300 and ∼30 lux, respectively. For the test, mice were introduced into the center zone of the elevated plus-maze and allowed to move freely for 10 minutes. All behaviors were recorded by a top-view infrared camera, and the time spent in each arm and the number of arm entries were measured and analyzed using EthoVision XT 10 software (Noldus).

<u>*5. Forced swim test*.</u> Mice were individually placed in a glass cylinder (15 × 30 cm) containing water (24°C ± 1°C; depth, 15 cm). All mice examined were forced to swim for 6 minutes, and the duration of immobility was recorded and measured during the final 4 minutes of the test. The latency to immobility from the start of the test (delay between the start of the test and appearance of the first bout of immobility, defined as a period of at least 1 second without any active escape behavior), and the duration of immobility (defined as the time not spent actively exploring the cylinder or trying to escape from it) were measured. Immobility time was defined as the time the mouse spent floating in the water without struggling, making only minor movements that were strictly necessary to maintain its head above water.

### Data analysis and statistics

All data are expressed as means ± SEM. All experiments were repeated using at least three independent cultures, and data were statistically evaluated using a Mann-Whitney *U* test, analysis of variance (ANOVA) followed by Tukey’s *post hoc* test, Kruskal-Wallis test (one-way ANOVA on ranks), paired two-tailed t-test (for electrophysiology experiments), or one-way ANOVA with Bonferroni’s *post hoc* test (for behavior experiments), as appropriate. Prism7 (GraphPad Software) was used for analysis of data and preparation of bar graphs. *P* values < 0.05 were considered statistically significant (individual *p* values are presented in figure legends).

