## Supplementary material for "The activity-dependent transcription factor Npas4 regulates IQSEC3 expression in somatostatin interneurons to mediate anxiety-like behavior": Manuscript

Kim et al.

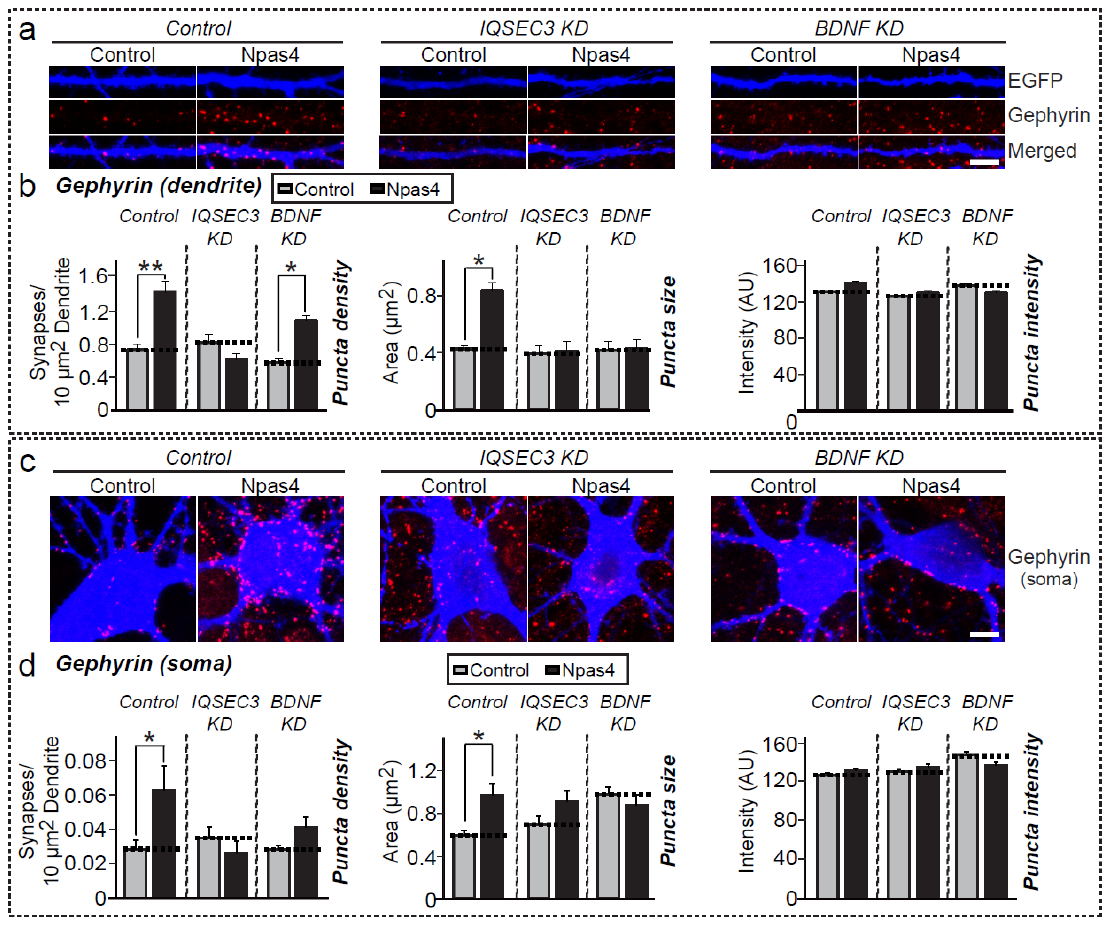
**Supplementary Figure 1.** IQSEC3 mediates GABAergic synapse development as an Npas4 target. (**a** and **c**) Representative images of cultured hippocampal neurons infected at DIV3 with lentiviruses expressing EGFP alone (Control), IQSEC3 KD or BDNF KD, and transfected at DIV10 with lentiviral constructs expressing EGFP alone (Control) or cotransfected with EGFP and Npas4 (Npas4). Dendrites (**a**) or cell bodies (**c**) of transfected neurons were analyzed by double-immunofluorescence labeling for gephyrin (red) and EGFP (blue) at DIV14. Scale bar, 10 μm (applies to all images). (**b** and **d**) Summary graphs of the effects of IQSEC3 KD or BDNF KD in Npas4-expressing neurons on gephyrin puncta density (left), gephyrin puncta size (middle), and gephyrin puncta intensity (right). Dendrites (**b**) and cell bodies (**d**) were quantified separately. For dendritic analyses, two or three dendrites per transfected neuron were chosen and group-averaged; n = 24 neurons/3 independent experiments. Data are presented as means ± SEMs (**p* < 0.05; ***p* < 0.01; non-parametric ANOVA with Kruskal-Wallis test, followed by *post hoc* Dunn’s multiple comparison test).

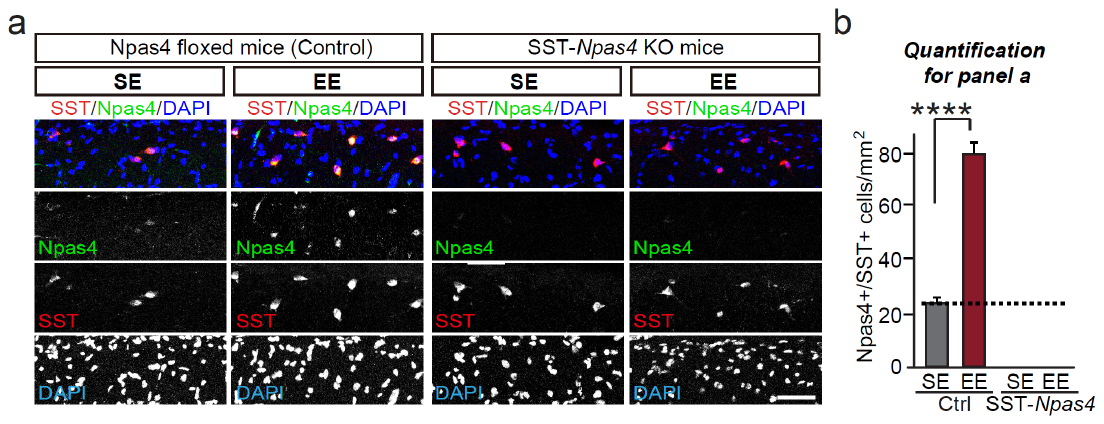

**Supplementary Figure 2.** Validation of EE-induced upregulation of Npas4 level using SST-*Npas4* KO mice. **(a)** Representative immunofluorescence images of the hippocampal CA1 SO layer from Control or SST-*Npas4* mice under SE or EE conditions, immunostained for Npas4 (green), SST (red), and DAPI (blue). Scale bar, 50 μm (applies to all images).

**(b)** Bar graphs summarizing the number of Npas4+ cells expressed in SST+ interneurons shown in (a). Data are presented as means ± SEMs (*****p* < 0.0001; non-parametric Mann-Whitney *U* test).

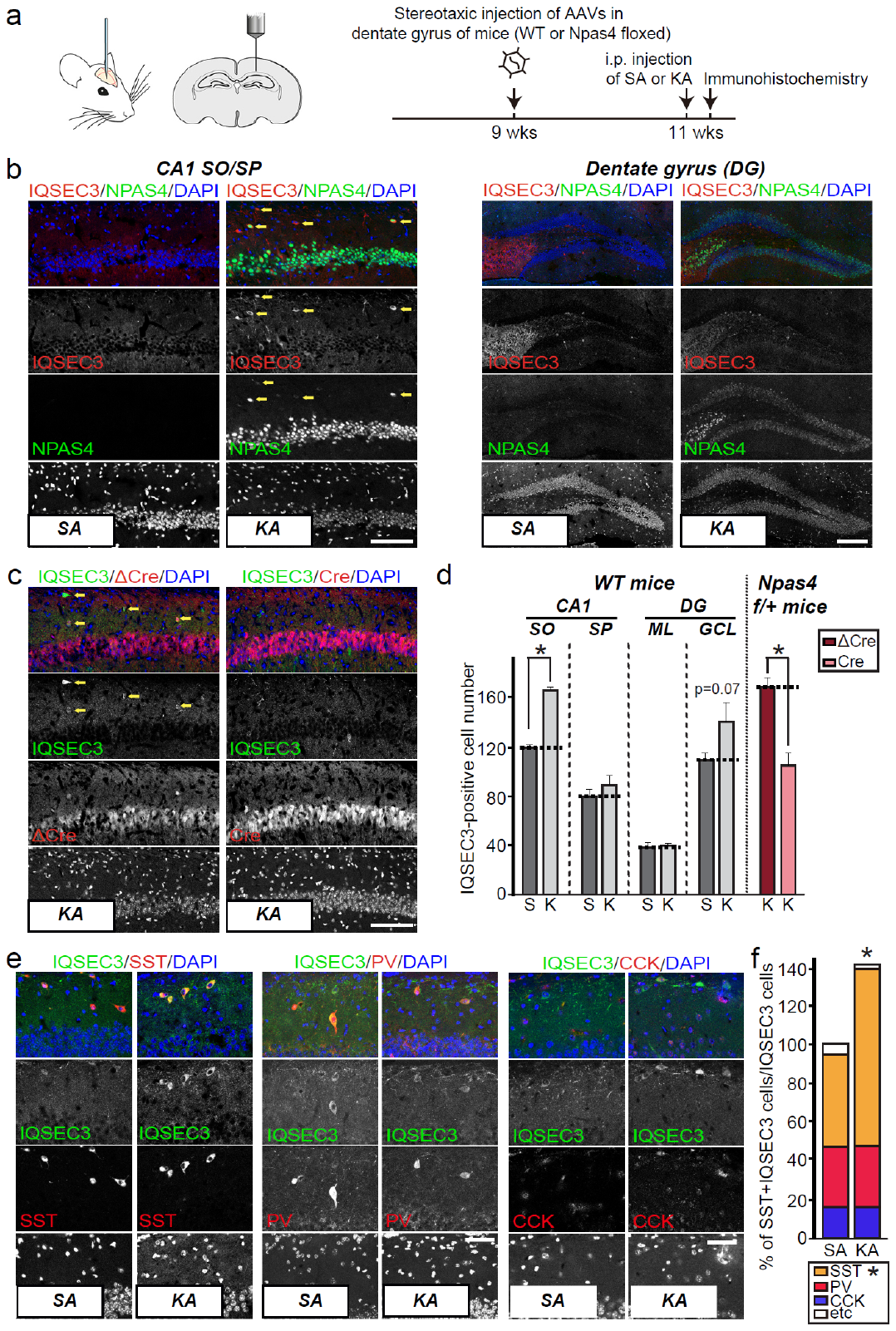

**Supplementary Figure 3.** IQSEC3 protein level is regulated by neuronal activity *in vivo****.***

**(a)** Strategy employed in immunohistochemistry experiments presented in panels (**b–f**).

**(b)** Representative immunofluorescence images of hippocampal CA1 (*left*) and DG (*right*) regions from 11-week-old adult WT mice intraperitoneally injected with either SA or KA, immunostained for IQSEC3 (red) and Npas4 (green), and counterstained with DAPI (blue). Yellow arrows denote neurons in which IQSEC3 was upregulated by injection of KA, but not SA. Scale bar, 100 μm (left; CA1 *SO/SP*) and 200 μm (right; *DG*). **(c)** Representative immunofluorescence images of hippocampal CA1 from Npas4^f/f^ mice injected with AAVs expressing mCherry fused to inactive (ΔCre) or active (Cre) Cre recombinase at ~9 weeks of age, intraperitoneally injected with KA when mice at ~11 week old, and stained for IQSEC3 (green), ΔCre or Cre (red), and counterstained with DAPI (blue). Scale bar, 100 μm (applies to all images). (**d)** Bar graphs summarizing quantitative results presented in (**b** and **c**). Abbreviations: DG, dentate gyrus; GCL, granule cell layer; K, kainic acid; ML, molecular layer; S, saline; SP, stratum pyramidale; SO, stratum oriens. Data are presented as means ± SEMs (**p* < 0.05 vs. individual saline controls; Mann-Whitney *U* test; n = 24 sections/4 mice for all conditions). **(e)** Representative immunofluorescence images of the hippocampal CA1 SO layer from 11-week old adult WT mice intraperitoneally injected with either SA or KA, immunostained for IQSEC3 (green), DAPI (blue), and SST (left), PV (middle), or CCK (right). Note that yellow arrows indicate the SST-positive interneurons with upregulated IQSEC3 levels. Scale bar, 50 μm (applies to all images). **(f)** Bar graphs summarizing quantitative results presented in (**e**). Data are presented as means ± SEMs (**p* < 0.05 vs. individual saline controls; Mann-Whitney *U* test; n = 18−24 sections/3−4 mice). Abbreviations: K, kainic acid; S, saline; SO, stratum oriens.

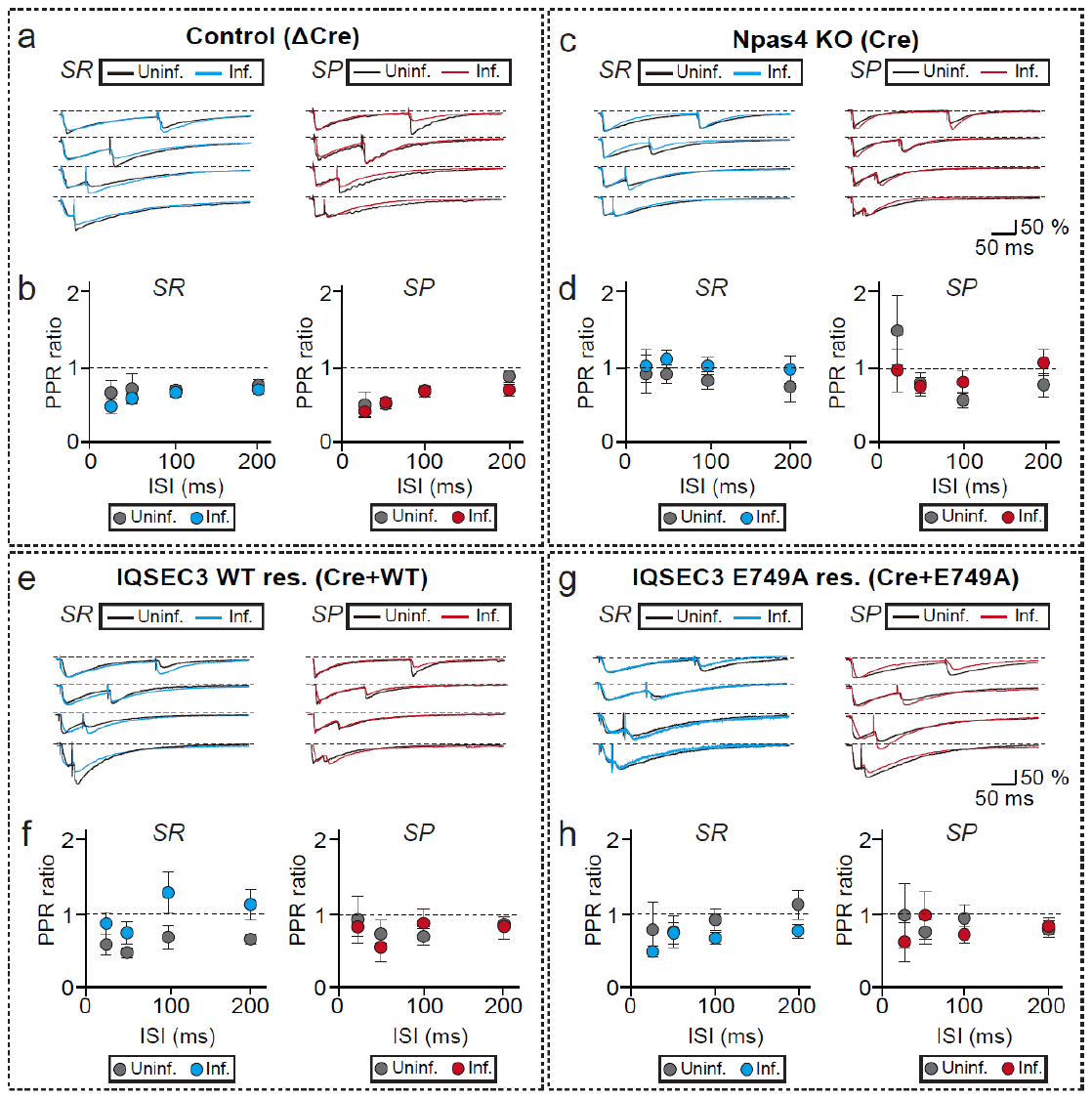

**Supplementary Figure 4.** No differences in PPRs of inhibitory postsynaptic currents between Npas4 WT and Npas4 KO neurons. Representative traces (**a**, **c**, **e**, **g**) and quantification (**b**, **d**, **f**, **h**) of IPSC-PPRs with the indicated interstimulus intervals (ISIs; 25–200 ms). Responses to stimulation of the SR (blue) or SP (red) layer were measured in neurons infected with AAV-ΔCre-EGFP, AAV-Cre-EGFP and AAV-IQSEC3-WT-T2A-tdTomato (IQSEC3 WT res.), or AAV-IQSEC3-E749A-T2A-tdTomato (IQSEC3 E749A res.), or in neighboring uninfected neurons (black), from mice housed under EE conditions. Data are presented as means ± SEMs (SR [blue]: Control (ΔCre), n = 7 pairs; Npas4 KO (Cre), n = 7 pairs; WT-rescue (Cre + IQSEC3 WT), n = 9 pairs; E749A-rescue (Cre + IQSEC3 E749A), n = 10 pairs; SP [red]: Control (ΔCre), n = 7 pairs; Npas4 KO (Cre), n = 7 pairs; WT-rescue (Cre + IQSEC3 WT), n = 8 pairs; and E749A-rescue (Cre + IQSEC3 E749A), n = 7 pairs). Scale bar, the percent change compared with uninfected neurons.

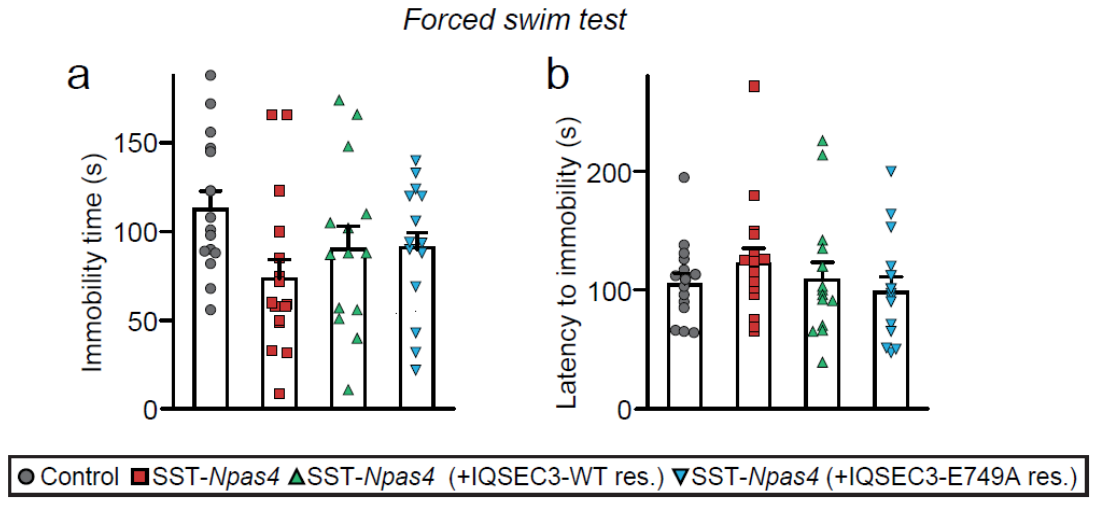

**Supplementary Figure 5.** No effect of Npas4 deletion in SST-expressing interneurons on depressive-like behavior. (**a** and **b**) Analysis of the depressive-like behavior by FS test in SST-*Npas4* mice injected with the indicated AAVs. Immobility time (**a**) and latency to immobility (**b**) were measured. Data are presented as means ± SEMs (Control, n = 10; SST-*Npas*4, n = 15; Rescue (+IQSEC3-WT), n = 14; and Rescue (+IQSEC3 E749A), n = 14; one-way ANOVA with Bonferroni’s *post hoc* test).

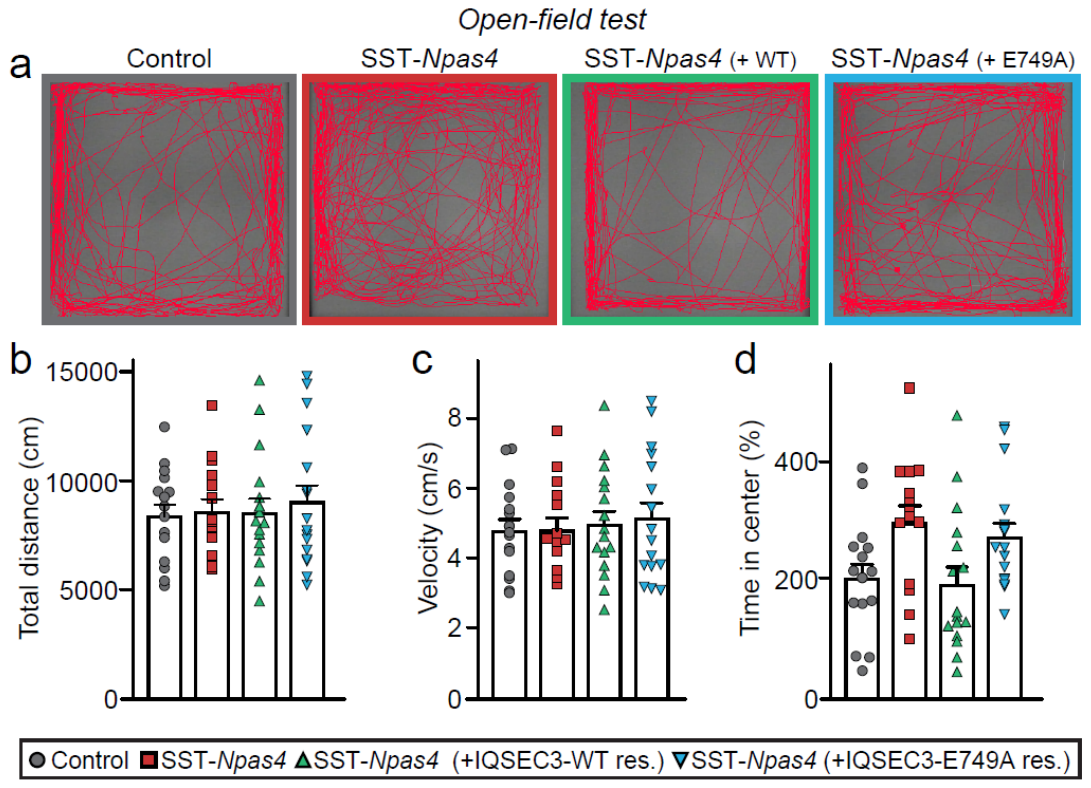

**Supplementary Figure 6.** No effect of Npas4 deletion in SST-expressing interneurons on locomotion. (**a–e**) Analysis of locomotor behavior by OF test in SST-*Npas4* mice injected with the indicated AAVs. Representative traces of locomotor activity (**a**) are presented. Total distance traversed (**b**), velocity (**c**), and time in the center zone (**d**) were measured. Data are presented as means ± SEMs (Control, n = 15; SST-*Npas*4, n = 14; Rescue (+IQSEC3-WT), n = 16; and Rescue (+IQSEC3 E749A), n = 16; one-way ANOVA with Bonferroni’s *post hoc* test).

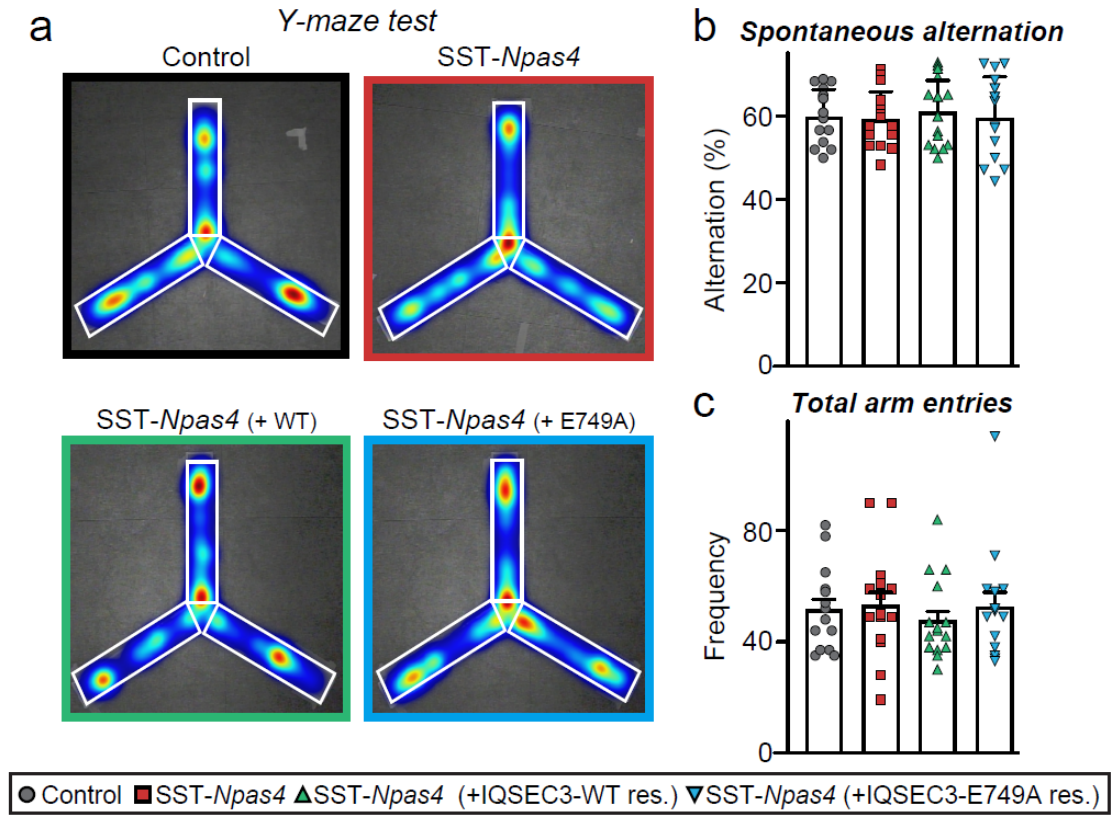

**Supplementary Figure 7.** No effect of Npas4 deletion in SST-expressing interneurons. (**a–c**) Analysis of the spatial working memory by Y-maze test in SST-*Npas4* mice injected with the indicated AAVs. Representative heat map showing movements of the indicated mice during the Y-maze test (**a**). Spontaneous alternation performance ratio (% of spontaneous alternations; **b**) and total number of arm entries (**c**) were measured. Data are presented as means ± SEMs (Control, n = 14; SST-*Npas*4, n = 15; Rescue (+IQSEC3-WT), n = 15; and Rescue (+IQSEC3 E749A), n = 14; one-way ANOVA with Bonferroni’s *post hoc* test).

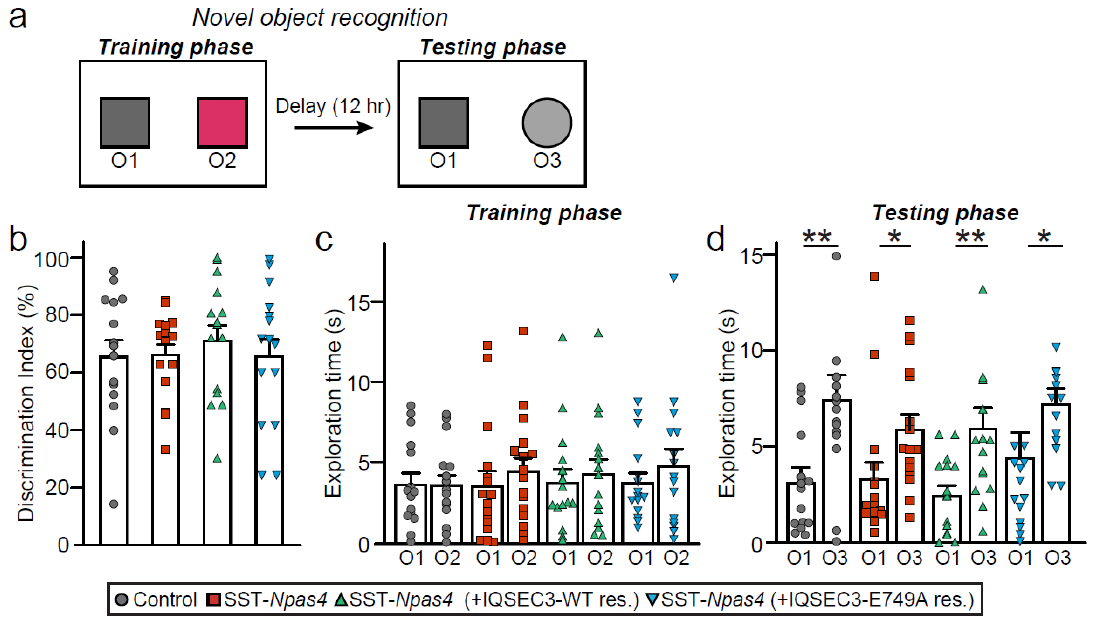

**Supplementary Figure 8.** No effect of Npas4 deletion in SST-expressing interneurons on novel object recognition memory. (**a–d**) Analysis of object recognition memory by NOR test in SST-*Npas4* mice injected with the indicated AAVs. Schematic diagram of the NOR test (**a**) is shown. Mice were allowed to explore two identical objects (denoted O1 and O2), and after a 12-hour delay, were exposed to two different objects: one familiar object from the training phase (O1) and one novel object (O3). Discrimination index (**b**) and exploration time for both training and testing phases (**c, d**) were measured. Data are presented as means ± SEMs (Control, n = 15; SST-*Npas*4, n = 16; Rescue (+IQSEC3-WT), n = 15; and Rescue (+IQSEC3 E749A), n = 15; **p* < 0.05 and *******p* < 0.01, one-way ANOVA with Bonferroni’s *post hoc* test).

| **Region** | **Forward Primer** | **Reverse Primer** |
| --- | --- | --- |
| **-10K** | CTATCACAGTAAATCTGCCT | GGTAGAATAGAATCATGGA |
| **promoter** | CTTCCTTCTTCCCTGGTAG | ACCTCTGGGCTTCGTGAG |
| **+14K** | GAGGAGCTGAGATGCAGTA | TCATGTTTGCCATTCAGATA |

**Supplementary Table 1.** Oligonucleotide sequences for ChIP analysis of the *Iqsec3* gene.

| **Experimental Group** | **Capacitance (pF)** | **Rise time (ms)** | **Decay time (ms)** |
| --- | --- | --- | --- |
| Control | 19.64 ± 1.19 | 0.98 ± 0.03 | 7.45 ± 0.13 |
| Npas4 KO | 19.29 ± 2.52 | 0.98 ± 0.04 | 7.26 ± 0.18 |
| WT rescue | 19.62 ± 1.60 | 1.03 ± 0.03 | 7.60 ± 0.09 |
| E749A rescue | 18.07 ± 1.06 | 1.03 ± 0.04 | 7.55 ± 0.12 |

**Supplementary Table 2.** Summary of the intrinsic electrophysiological properties of Npas4 WT and Npas4 KO hippocampal somatostatin-expressing neurons.

| **Experimental Group** | | **Capacitance (pF)** | **Input Resistance (MΩ)** | **Series Resistance (MΩ)** |
| --- | --- | --- | --- | --- |
| Control (ΔCre) | Uninfected | 118.77 ± 12.37 | 178.78 ± 16.10 | 27.45 ± 1.11 |
|  | Infected | 113.99 ± 18.20 | 181.93 ± 14.67 | 26.84 ± 0.88 |
| Npas4 KO (Cre) | Uninfected | 103.33 ± 14.62 | 277.04 ± 35.02 | 26.28 ± 1.25 |
|  | Infected | 134.74 ± 17.79 | 193.99 ± 43.20 | 27.26 ± 0.48 |
| WT rescue | Uninfected | 129.25 ± 18.14 | 297.59 ± 75.96 | 25.33 ± 1.64 |
|  | Infected | 130.30 ± 15.49 | 193.90 ± 26.01 | 25.18 ± 1.16 |
| E749A rescue | Uninfected | 104.81 ± 9.52 | 209.03 ± 34.27 | 23.68 ± 1.05 |
|  | Infected | 133.08 ± 17.11 | 207.57 ± 39.03 | 24.82 ± 1.03 |

**Supplementary Table 3.** Summary of the intrinsic electrophysiological properties of Npas4 WT and Npas4 KO hippocampal CA1 pyramidal neurons.
